# Differential Regulation of Excitatory vs Inhibitory Synaptic Release by Complexin/CPX-1 and CAPS/UNC-31 at the *C. elegans* Neuromuscular Junctions

**DOI:** 10.1101/2023.08.25.554737

**Authors:** Ya Wang, Chun Hin Chow, Mengjia Huang, Randa Higazy, Neeraja Ramakrishnan, Cuntai Zhang, Shuzo Sugita, Shangbang Gao

**Author notes:** **Equal contributions** (Ya Wang, Chun Hin Chow). **Correspondence**: Shuzo Sugita, Shangbang Gao.

## Abstract

The excitation and inhibition (E/I) balance at neuromuscular junctions plays a crucial role in coordinating animal motor behavior. Prominent synaptic vesicle secretory regulatory proteins, specifically complexin and CAPS (Calcium-dependent Activator Protein for Secretion), have not garnered sufficient attention for E/I balance regulation. Here, we investigate the roles of complexin/CPX-1 and CAPS/UNC-31 in excitatory vs inhibitory synapses of *C. elegans* neuromuscular junctions. In our study, *cpx-1* null mutants displayed remarkable reduction evoked release in both excitatory and inhibitory synapses. Intriguingly, these mutants exhibited an enhanced level of spontaneous release, particularly within the context of excitatory synapse. This enhancement aligns with its “clamp” role that preventing SV from fusing with the presynaptic membrane. Additionally, a clamping-specific knockin mutant *cpx-1(Δ12)*, which displayed no alterations in evoked release at either type of synapse, also revealed a biased substantial increase in excitatory spontaneous release. In contrast, *unc-31* null mutation, with normal spontaneous release, led to a more pronounced decreased evoked release in the excitatory synapse independent of dense-core vesicle regulation. Intriguingly, we found that the enhanced excitatory spontaneous release observed in *cpx-1* mutants was abolished by *unc-31* in a Ca^2+^-dependent manner, implying that UNC-31 is essential for maintaining a high level of spontaneous fusion events. Collectively, our findings unveil a distinct regulatory pattern governing excitatory and inhibitory synaptic release, orchestrated by the interplay of CPX-1 and UNC-31. Notably, we uncover an unforeseen role of UNC-31 in influencing CPX-1’s “clamp” function, further adding complexity to the neural dynamics, which may underlie complex behavioral phenotypes observed in *C. elegans*.

## Introduction

Maintaining excitatory and inhibitory (E/I) balance is a fundamental property of functional neuronal circuits. The balance between excitatory and inhibitory synaptic transmission in the motor circuitry is critical for generating coordinated locomotion. Altered E/I balance in the brain underlies epilepsy and is implicated in psychiatric disorders such as Autism (Sohal & Rubenstein, 2019; van van Hugte et al., 2023; Yizhar et al., 2011). Modulation of synaptic vesicular release of neurotransmitters is one important mechanism for controlling the relative strength/weight of excitatory and inhibitory inputs (Deng et al., 2021). In both types of synapses, the neuronal <u>S</u>oluble <u>N</u>–ethylmaleimide sensitive factor <u>A</u>ttachment protein <u>RE</u>ceptor (SNARE) proteins drive the fusion of neurotransmitter-filled synaptic vesicles with the plasma membrane (Söllner et al., 1993). Much of our previous understanding of the regulation of SNARE-mediated vesicular fusion comes from mammalian, glutamatergic excitatory synapses. However, the regulatory mechanisms may differ between excitatory and inhibitory synapses, yet such differential regulation is largely unknown, at least in *C. elegans*.

The highly regulated and synchronous process of vesicular fusion at the synapse requires an array of regulatory proteins at the presynaptic terminal. In most synapses that have been studied, a pool of primed (i.e. readily released) vesicles is maintained until synaptotagmin senses the Ca^2+^ influx (Martens & McMahon, 2008; Rizo & Rosenmund, 2008). Complexin is a key synaptic vesicle secretory regulatory protein that functionally and structurally interacts with synaptotagmin to keep vesicles at primed states before evoked release (Ramakrishnan et al., 2020; Tokumaru et al., 2008; Zhou et al., 2017). Complexins are evolutionarily conserved proteins composed of N-terminal and C-terminal regions with a central α helix and an accessory α helix (Chen et al., 2002; McMahon et al., 1995). Studies on mice, *Drosophila* and *C. elegans* complexin null mutants showed consistent reductions in synchronous Ca^2+^-dependent evoked excitatory exocytosis (Chang et al., 2015; Hobson et al., 2011; Huntwork & Littleton, 2007; Jorquera et al., 2012; Martin et al., 2011; Strenzke et al., 2009; Xue et al., 2010; Xue et al., 2008). The N-terminal domain is critical for promoting fast fusion events (Hao et al., 2023; Maximov et al., 2009). On the other hand, the C-terminal domain (CTD) of complexin is proposed to perform the inhibitory clamping function, where synaptic vesicles are “clamped” at the primed state (Hao et al., 2023; Iyer et al., 2013; Kaeser-Woo et al., 2012; Maximov et al., 2009; Ramakrishnan et al., 2020; Rizo, 2022). The inhibitory function of complexin is evident in which with a decreased or fully abolished complexin expression, spontaneous neurotransmitter release is increased (Giraudo et al., 2006; Hobson et al., 2011; Jorquera et al., 2012; Schaub et al., 2006; Tang et al., 2006). However, this idea remains controversial since a few mammalian studies show reduced spontaneous release in the absence of complexin (Chang et al., 2015; Strenzke et al., 2009; Xue et al., 2010; Xue et al., 2008). Regardless, the dual function of complexin in promoting synchronous release and inhibiting asynchronous release is largely based on excitatory synapses. Removing all three complexin isoforms in cultured GABAergic striatal neurons was reported to decrease both evoked and spontaneous releases (Xue et al., 2008), in contrast with the dual regulation in many excitatory synapses. In *C. elegans*, expression of complexin in excitatory acetylcholine motor neurons, but not in inhibitory GABAergic motor neurons, rescued the aldicarb hypersensitivity and impaired locomotion speed in *cpx-1* mutants (Hobson et al., 2011; Martin et al., 2011). Thus, the role of complexins at the excitatory and inhibitory synapses need to be independently studied. Furthermore, in mice cortical neurons, complexin was indicated to clamp spontaneous exocytosis by blocking a secondary Ca^2+^ sensor, in addition to synaptotagmin-1 (Yang et al., 2010). However, the predicated presynaptic partner interacting with complexin remain uncovered.

CAPS is another crucial synaptic vesicle secretory regulatory protein (Charlie et al., 2006), primarily recognized for its involvement in the modulation of dense-core vesicle (DCV) exocytosis (Bastian et al., 2020; Berwin et al., 1998; Lin et al., 2010; Olsen et al., 2003; Rupnik et al., 2000; Tandon et al., 1998; Zhou et al., 2007). In contrast to UNC-13, which functions as a critical priming factor, UNC-31’s role in regulating synaptic vesicle exocytosis has been a topic of extensive debate. As an evolutionarily conserved calcium-binding protein, CAPS/UNC-31 contains two major domains: a central PH domain that binds to phospholipids and a C-terminal domain that is homologous to Munc13’s protein domains involved in synaptic vesicle priming. Null mutants for homologs of CAPS in *C. elegans (unc-31)* and *Drosophila* (*dCAPS*) show a loss of secretion in neurotransmitters and neuropeptides (Ann et al., 1997; Avery et al., 1993). Speese et al. further showed that *unc-31 null C. elegans* mutants exhibited reduced peptide release *in vivo* but had normal stimulated synaptic vesicle recycling in cultured neurons (Speese et al., 2007). With the homologous domain to Munc13, CAPS/UNC-31 is proposed to be docking DCVs (Imig et al., 2014; Zhou et al., 2019). However, several lines of evidence suggest synaptic transmission is also, at least in part, dependent on CAPS/UNC-31. Renden et al. showed that dCAPS mutants exhibit mild reductions in synaptic transmission (Renden et al., 2001). CAPS1 and CAPS1&2 double knockout mice exhibit a strong decrease in synaptic vesicle exocytosis at excitatory synapses (Jockusch et al., 2007). Importantly, loss of UNC-31 in *C. elegans* also revealed a decreased evoked excitatory transmission (Cornell et al., 2022; Gracheva et al., 2007). Nevertheless, a reduction in synaptic transmission by the deletion of CAPS/UNC-31 is generally considered secondary to a reduction in DCV release in the *C. elegans* field (Speese et al., 2007). Several questions regarding CAPS/UNC-31 are crucial: 1) is there a differential regulatory effect by CAPS/UNC-31 on excitatory vs inhibitory synapses, 2) whether CAPS/UNC-31, sharing Munc13’s priming domains, directly controls the release of synaptic vesicles independent of its docking function of DCV, and 3) how CAPS/UNC-31 functionally interacts with other synaptic regulatory proteins such as complexin.

In this study, we investigate the roles of Complexin/CPX-1, CAPS/UNC-31 and their interaction in synaptic vesicle exocytosis at both excitatory and inhibitory synapses using the *C. elegans* model. Our results reveal that while *cpx-1* null mutation decreases evoked release in both excitatory and inhibitory synapses, it enhances spontaneous release primarily in the excitatory synapses. We show that CAPS/UNC-31 plays a significant role in evoked, but not spontaneous, synaptic vesicle exocytosis independent of dense-core vesicle regulation. Deficits in evoked exocytosis of *unc-31* are stronger in the excitatory synapses compared to the inhibitory ones. Intriguingly, when spontaneous release is enhanced by the removal of complexin/CPX-1 clamping, CAPS/UNC-31 becomes indispensable for the maintenance of enhanced spontaneous exocytosis and this process can only be seen in the presence of extracellular calcium. Our results demonstrate the differential regulation of excitatory vs inhibitory synaptic vesicle exocytosis by complexin/CPX-1 and CAPS/UNC-31. The requirement of UNC-31 for CPX-1’s inhibitory function may underlie the basis of complex behavioral phenotypes of *cpx-1* and *unc-31* mutants.

## Results

### *cpx-1* null mutation enhances the frequency and amplitude of miniature PSCs, but not inhibitory miniature PSCs, while it decreased both excitatory and inhibitory evoked PSCs

Most *C. elegans* NMJ electrophysiological analyses on synaptic transmission have been limited to excitatory transmission, leaving the mechanism of inhibitory synaptic release less well understood. We examined the roles of two presynaptic regulatory proteins of exocytosis, complexin/CPX-1 and CAPS/UNC-31, in both excitatory and inhibitory neuromuscular junctions (NMJs). The selective expression of channel rhodopsin at excitatory cholinergic or inhibitory GABAergic neurons enables reliable measurement of exocytosis underlying neurotransmission in *C. elegans*. We used *cpx-1(ok1552)*, a *cpx-1 null* mutant, or *unc-31(e928)*, an *unc-31 null* mutant, with *zxIs6* (excitatory synapse) or *zxIs3* (inhibitor synapse) lines to measure exocytosis using optogenetics (Liewald et al., 2008).

*cpx-1 null* mutants exhibited enhanced spontaneous release and reduced evoked release from excitatory motor neurons (Hobson et al., 2011; Martin et al., 2011). Given that *C. elegans* neuromuscular junctions (NMJs) receive both cholinergic and GABAergic inputs simultaneously, mixed excitatory (predominantly) and inhibitory (minor) miniature post-synaptic currents (mPSCs) are collected with the standard electrophysiological recording condition (Richmond & Jorgensen, 1999, Methods). Consistent with previous observations (Hobson et al., 2011; Martin et al., 2011), we found that *cpx-1(ok1552)* increases the frequency of the mPSCs, compared to wild-type animals (**Fig. 1A, B**). Surprisingly, in contrast to all previously studied *cpx-1* mutants that exhibited no discernible effect on mPSC amplitude, the *cpx-1(ok1552)* mutant displays a slight but significant increase in mPSC amplitude (P< 0.01, **Fig. 1C**). On the other hand, excitatory evoked post-synaptic current (eEPSC) of *cpx-1(ok1552)* is drastically decreased compared to the wild type (**Fig. 1D-F**). Indeed, the role of *C. elegans* CPX-1 is in line with the widely accepted perspective of its dual impact – facilitating evoked release while exerting an inhibitory influence on spontaneous synaptic transmission.

**Figure 1.**
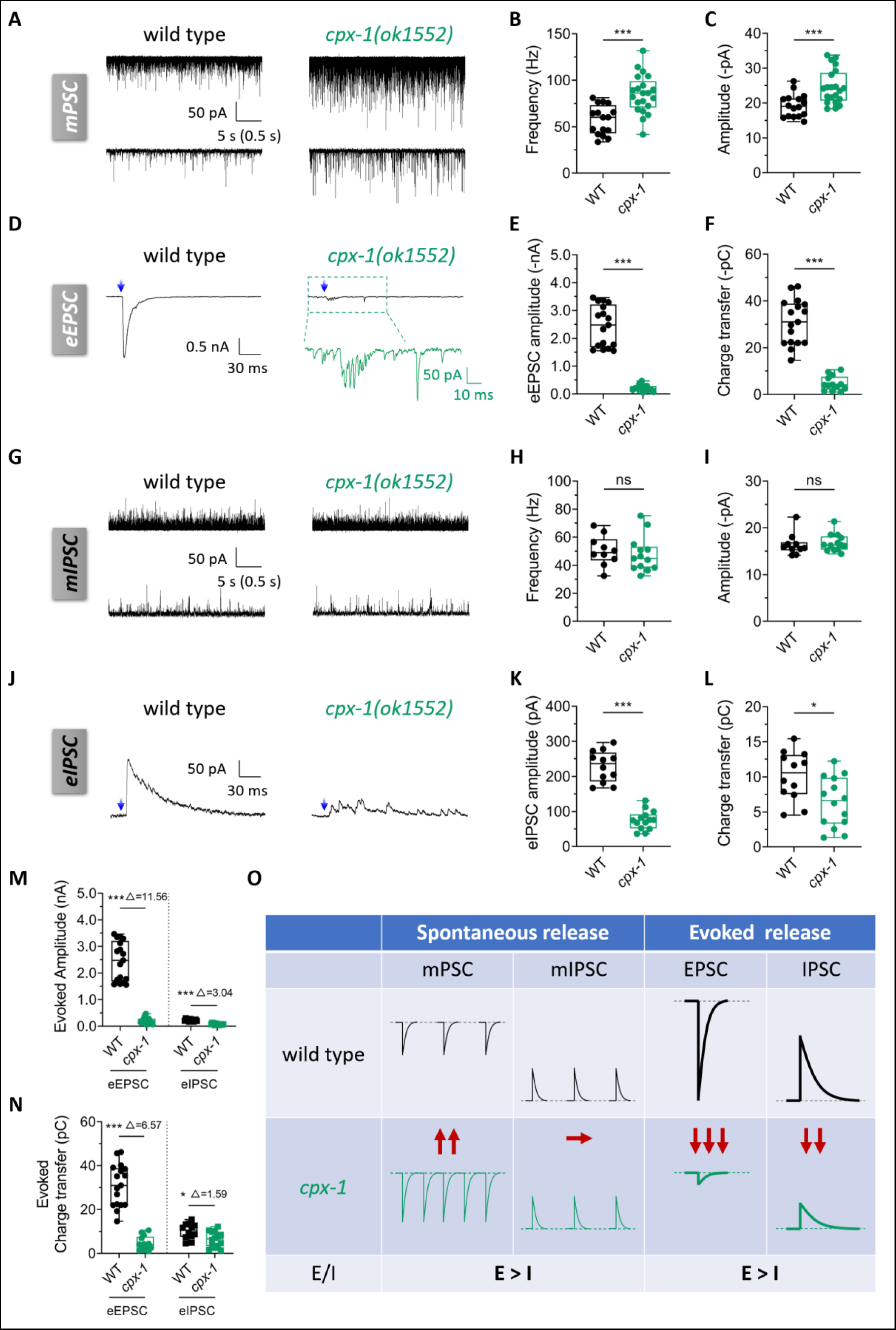
Enhanced excitatory spontaneous synaptic vesicle release in cpx-1(ok1552) mutants at C. elegans NMJ. (A) Representative traces of miniature postsynaptic currents (mPSCs) recorded from N2 wild-type (left) and *cpx-1(ok1552)* (right) worms, shown at different timescales. (B and C) Quantification of the mPSCs frequency (B) and amplitude (C) in wild-type and *cpx-1(ok1552)* worms. (D) Representative traces of evoked excitatory postsynaptic currents (eEPSCs) recorded from WT (left) and *cpx-1(ok1552)* (right) worms in the *zxIs6* [P*unc-17*::ChR2::YFP + *lin-15*(+)] transgenic background with 10 ms blue light illumination (blue arrow). The lower trace of *cpx-1(ok1552)* (green) shows a zoomed-in region from the upper trace. (E and F) Quantification of the EPSCs amplitude (E) and charge transfer (F) in WT and *cpx-1(ok1552)* worms. Recordings were made from body wall muscle cells at a holding potential of −60 mV. (G) Representative traces of mIPSCs recorded from WT (left) and *cpx-1(ok1552)* (right) worms, displayed at different timescales. (H and I) Quantification of the mIPSC frequency (H) and amplitude (I) in WT and *cpx-1(ok1552)* worms. (J) Representative traces of inhibitory postsynaptic currents (eIPSCs) recorded from WT (left) and *cpx-1(ok1552)* (right) worms in the *zxIs3* [P*unc-47*::ChR2::YFP + *lin-15*(+)] transgenic background with 10 ms blue light illumination (blue arrow). (K and L) Quantification of the eIPSCs amplitude (K) and charge transfer (L) in WT and *cpx-1(ok1552)* worms. Muscles were held at −10 mV with 0.5 mM ionotropic acetylcholine receptor blocker D-tubocurarine (d-TBC). (M and N) Box-and-whisker plots of the eEPSC (left) and eIPSC (right) amplitude (M) and charge transfer (N) in WT and *cpx-1(ok1552)* worms. The numbers after △ represent the ratio of the average values of WT and *cpx-1(ok1552)*.(O) Schematic representation illustrating the differential changes in excitatory and inhibitory synaptic vesicle release in *cpx-1* mutant. Data are presented as box-and-whisker plots, with the median (central line), 25th–75th percentile (bounds of the box), and 5th–95th percentile (whiskers) indicated. Student’s t-test was performed, *p<0.05; **p<0.01; ***p<0.001; ns, not significant. Error bars represent standard error of the mean (SEM).

These mPSCs comprise elements from both excitatory and inhibitory minis-components. Whether CPX-1 serves equal roles in excitatory and inhibitory synapses remains unknown. To clarify this, we specifically recorded GABAergic transmission in *cpx-1(ok1552)*. To do so, we employed D-tubocurarine (d-TBC, 0.5 mM), a broad-spectrum blocker of *C. elegans* ionotropic AChRs, to isolate the miniature inhibitory postsynaptic currents (mIPSCs) (Maro et al., 2015; Richmond & Jorgensen, 1999). However, in comparison with the wild type, neither the frequency nor amplitude of mIPSCs was altered in *cpx-1(ok1552)* (**Fig. 1G-I**). The observation that CPX-1 is unlikely to be necessary for mIPSCs in inhibitory synapses suggests that the increased mPSCs observed in *cpx-1* mutants may be attributed to its function in excitatory synapses. Nevertheless, evoked inhibitory postsynaptic currents (eIPSCs) is also decreased in *cpx-1(ok1552)* mutant (**Fig. 1J-L**). Thus, unlike the case for excitatory NMJs, decreased evoked release is not coupled with enhanced spontaneous release in inhibitory synapses that lack *cpx-1*. Considering that most components of the release machinery, including complexins, participate in both excitatory and inhibitory synapses, the specific disruption of CPX-1’s function in excitatory but not inhibitory spontaneous release suggests a coordinated or balanced mechanism between CPX-1 and other release machinery (see discussion).

To delve deeper into the distinct impacts of *cpx-1(ok1552)* on excitatory and inhibitory transmission, we conducted a comparative analysis of its preferential influence on excitatory synapses versus inhibitory synapses. While mIPSC is unaltered in *cpx-1(ok1552)* mutant animals, mPSC frequency and amplitude are increased (**Fig 1A-F**). Thus, spontaneous release at the NMJ without CPX-1 might be driven to a heightened excitatory state (**Fig 1O**, *left*). For evoked release, we analyzed the relative magnitudes (Δ = I_WT_ / I*_cpx-1_*) and observed that in *cpx-1* mutants, the peak amplitude of eEPSC decreased by a factor of Δ=11.56, while the reduction in eIPSC was only Δ=3.04 (**Fig 1M**). Consequently, this yielded a notable 3.8-fold disparity in the relative contributions of CPX-1 to excitation to inhibition. Similar trend was observed when we compared the relative change in transferred charges, where showed a 4.1-fold disparity between eEPSC (Δ = 6.57) and eIPSC (Δ = 1.59) (**Fig 1N**). As such, evoked transmission in *cpx-1(ok1552)* mutants preferentially weaken excitatory strength compared to the inhibition (**Fig 1O**, *right*).

Collectively, these findings imply that complexin/CPX-1 exerts a more pronounced influence on excitatory synapses compared to inhibitory synapses (E > I) at the *C. elegans* neuromuscular junction.

### A clamping-defective *cpx-1* mutant exhibit more prominent defect at the excitatory synapse

Decreased synchronous release observed in *cpx-1* has been considered in part due to the enhanced spontaneous release depleting the primed vesicles. In *C. elegans*, no clamping-specific mutant that preserves evoked release has been identified thus far. Therefore, we generated a clamping-specific mutant in this study. The CTD of CPX-1 is critical for its suppression of spontaneous release or clamping function (Hao et al., 2023; Iyer et al., 2013; Kaeser-Woo et al., 2012; Maximov et al., 2009). In addition, Wragg et al. showed that deletion of the last 12 residues or exchanging L117 and V121 with glutamate (L117E,V121E, or LV/EE) point mutations of CPX-1 led to the loss of binding ability to liposomes (Wragg et al., 2013). Based on the functional and biochemical studies, we generated two knock-in (KI) lines using CRISPR/Cas9 (Methods). First is the deletion of C-terminal 12 residues (Δ12) and the second is the LV/EE mutant (**Fig 2A**). We explored the clamping functions of the KI mutants in excitatory and inhibitory synapses in vivo and compared their phenotypes with the null mutant, *cpx-1(ok1552)*.

**Figure 2.**
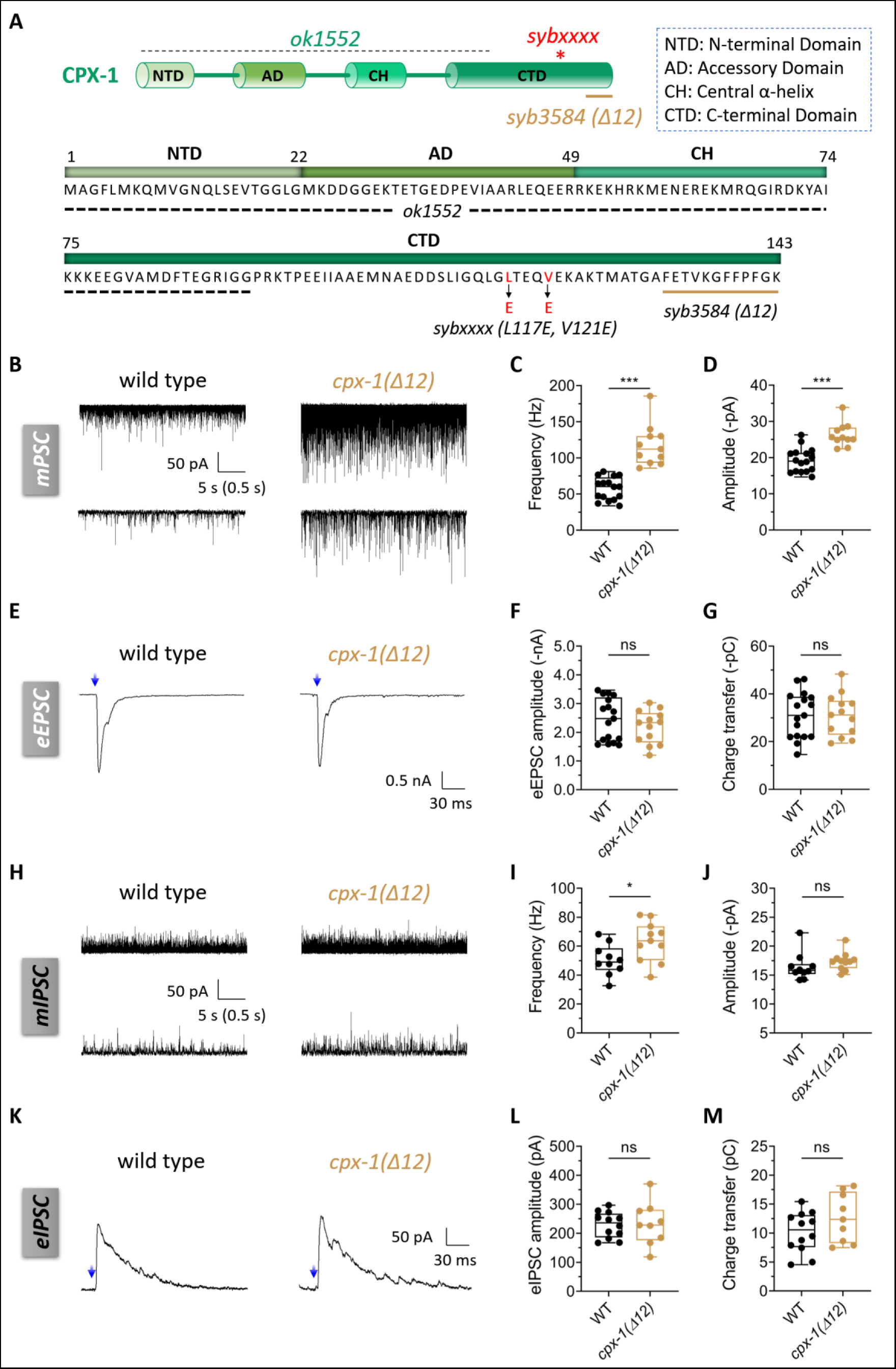
Enhancement of excitatory spontaneous release by cpx-1(Δ12) mutation without impact on evoked release. (A) Cartoon representation and amino acid sequence of CPX-1, indicating the N-terminal domain (NTD), accessory domain (AD), central α-helix (CH) and C-terminal domain (CTD). Mutation sites of *ok1552* (dash line), *syb3584* (brown line) and *syb3665 (red stars)* are labeled respectively. (B) Representative traces of mPSCs recorded from WT and *cpx-1(syb3584)*, which have a truncation of the last 12 amino acids in the CTD, referred to as *cpx-1(Δ12)*. (C and D) Quantification of the mPSCs frequency (C) and amplitude (D) in WT and *cpx-1(Δ12)* worms. (E-G) Representative traces and quantification of evoked EPSCs recorded from WT and *cpx-1(Δ12)* worms. (H-J) Representative traces and quantification of mIPSCs recorded from WT and *cpx-1(Δ12)* worms, displayed at different timescales. (K-M) Representative traces and quantification of evoked IPSCs recorded from WT and *cpx-1(Δ12)*. Data are presented as box-and-whisker plots, with the median (central line), 25th–75th percentile (bounds of the box), and 5th–95th percentile (whiskers) indicated. Student’s t-test was performed, *p<0.05; ***p<0.001; ns, not significant.

We utilized two assays to behaviorally evaluate synaptic transmission. Thrashing, which is the motility measurement of worms, indicates the strength of evoked EPSC and indirectly reflects E/I balance in generating sinusoidal movement. Another assay is measuring aldicarb sensitivity, where aldicarb is an acetylcholinesterase inhibitor preventing the removal of acetylcholine at the synapse. Increased aldicarb sensitivity is commonly associated with heightened excitatory transmission in *C. elegans*, but it could also stem from a disruption in the E/I balance. Consistent with prior findings (Hobson et al., 2011), *cpx-1(ok1552)* null mutant exhibits reduced thrashing and enhanced aldicarb sensitivity (**Fig S1A, B**). Reduced thrashing is consistent with the decrease in eEPSC (**Fig 1D-F**) while increased aldicarb sensitivity is explained by the increase in mPSC (**Fig 1A-C**). For the CTD KI mutants, *cpx-1(Δ12, syb3584)* showed better motility and more dramatic enhancement in aldicarb sensitivity compared to *cpx-1(ok1552)* (**Fig S1A, B**). On the other hand, *cpx-1(LV/EE, syb3665)* exhibits a similar movement level and aldicarb sensitivity to *cpx-1 null* (**Fig S1A, B**).

*cpx-1(Δ12)* retains better movement than *cpx-1 null* and shows further heightened aldicarb sensitivity; both phenotypes are consistent with the genetic lesion selectively disrupting CPX-1’s clamping ability without altering evoked synaptic transmission. Indeed, measuring synaptic transmission with electrophysiology, we found that *cpx-1(Δ12)* retains normal evoked release in both excitatory (**Fig 2E-G**) and inhibitory synapses (**Fig 2K-M**). As expected with the high aldicarb sensitivity, *cpx-1(Δ12)* strongly increases the frequency and amplitude of mPSC (**Fig 2B-D**). Surprisingly, different to no effect of *cpx-1* null mutant on mIPSC, *cpx-1(Δ12)* also slightly increases the mIPSC frequency with unaltered amplitude (**Fig 2H-J**). Given that evoked release is normal in the clamping-specific mutant *cpx-1(Δ12)*, we showed that evoked and spontaneous releases can be segregated where enhanced spontaneous activity does not necessarily deplete primed vesicles for synchronous release. Comparing the relative magnitudes of frequency (Δ = I_WT_ / I*_cpx-1_*) in spontaneous activities, the increase in mIPSC is weaker than that in the excitatory synapses (**Fig S2A-E**). Thus, in spontaneous release, *cpx-1(Δ12)* also displayed altered E/I balance with a dominating excitatory transmission (**Fig S2E**).

### *unc-31* null mutation decreases evoked exocytosis more strongly in the excitatory synapse compared to inhibitory synapse

Our analyses of CPX-1 demonstrated unexpected differences in its regulation of excitatory vs inhibitory synaptic release. we decided to investigate other regulatory partners in the two types of synapses. In the presynaptic secretion machinery, prominent proteins, such as Munc-13, Munc-18, and Synaptotagmins, have been extensively studied. Previous studies in *unc-31* null showed contradictory results in excitatory evoked release whereas its phenotype in the inhibitory synapse has not been investigated (Speese et al., 2007;Jockusch et al., 2007;Cornell et al., 2022; Gracheva et al., 2007). Here, we investigated the phenotype of *unc-31(e928)* null mutant.

We found that *unc-31(e928)* does not exert an impact on spontaneous release in either excitatory or inhibitory synapses, as evidenced by the unchanged frequency and amplitude of mPSCs and mIPSCs (**Fig 3A-C, G-I**). However, it decreases both eEPSC and eIPSC (**Fig 3D-F, J-L**). These findings diverge from the observations made by Speese et al., who noted no alterations in excitatory evoked release (Speese et al. 2007), but consistent with Gracheva et al. showing a reduction in eEPSC (Gracheva et al., 2007). Interestingly, the relative decreases in eEPSC amplitude and charge transfer are larger than in eIPSC by analyzing (Δ = I_WT_ / I*_unc-31_*) (**Fig 3M-N**). Therefore, *unc-31(e928)* tilts the E/I balance to a stronger inhibition side (**Fig 3O**). This can potentially explain the phenotype of *unc-31(e928)* mutants which show a rigid and straight posture indicating a stronger drive to muscle relaxation (Avery et al., 1993). Thus, similar to CPX-1, loss of UNC-31 induces different phenotypes in the two types of synapses, and there is more regulation of excitatory synapses.

**Figure 3.**
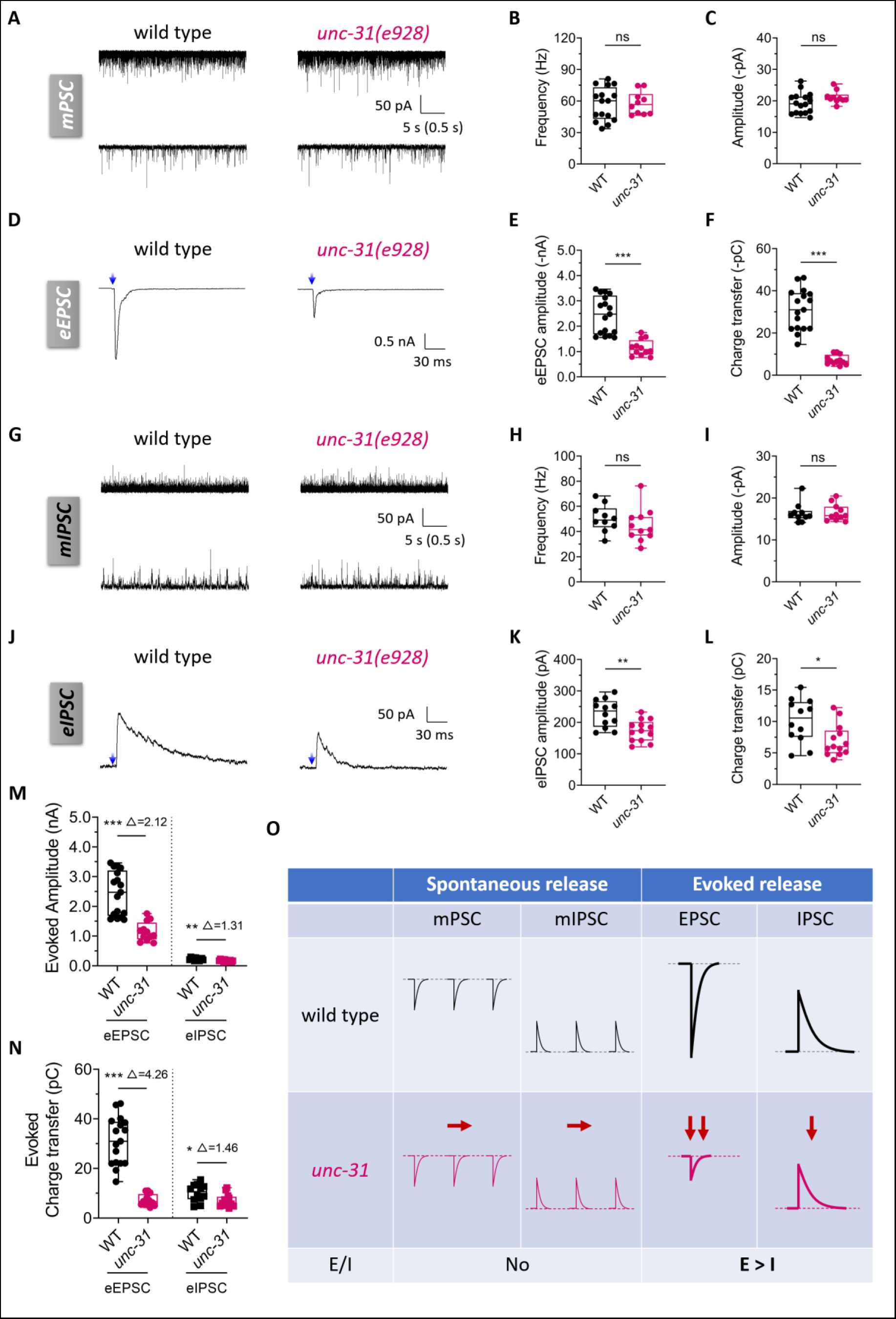
UNC-31/CAPS differentially regulates evoked cholinergic and GABAergic neurotransmission. (A-C) Representative traces and quantification of mPSCs recorded from WT and *unc-31(e928)* worms. (D) Representative traces of evoked EPSCs recorded from WT (left) and *cpx-1(ok1552)* (right) worms in the *zxIs6* [P*unc-17*::ChR2::YFP + *lin-15*(+)] transgenic background with 10 ms blue light illumination (blue arrow). (E and F) Quantification of the eEPSCs amplitude (E) and charge transfer (F) from WT and *cpx-1(ok1552)* worms. (G-I) Representative traces and quantification of mIPSCs recorded from WT (left) and *cpx-1(ok1552)* (right) worms, displayed at different timescales. (J) Representative traces of evoked IPSCs recorded from WT (left) and *cpx-1(ok1552)* (right) worms in the *zxIs3* [P*unc-47*::ChR2::YFP + *lin-15*(+)] transgenic background with 10 ms blue light illumination (blue arrow). (K and L) Quantification of the eIPSCs amplitude (K) and charge transfer (L) from WT and *cpx-1(ok1552)* worms. Muscles were held at −10 mV with 0.5 mM d-TBC. (M and N) Box-and-whisker plots of the amplitude (M) and charge transfer (N) of the EPSC (left)/IPSCs (right) in WT and *unc-31(e928)* worms. The numbers after △ represent the ratio of the average values of WT and *unc-31(e928)*. (O) Schematic representation illustrating the differential changes in excitatory and inhibitory synaptic vesicle release of *unc-31* mutant. Data are presented as box-and-whisker plots, with the median (central line), 25th–75th percentile (bounds of the box), and 5th–95th percentile (whiskers) indicated. Student’s t-test was performed, *p<0.05; **p<0.01; ***p<0.001; ns, not significant.

### Regulation of synaptic vesicle exocytosis by UNC-31 is independent of dense-core vesicle-mediated neural transmission

Alteration in synaptic transmission observed by the deletion of CAPS/UNC-31 has been considered secondary to a dysregulation of DCV release at least in the *C. elegans* field (Charlie et al., 2006; Speese et al., 2007). However, the differential reduction in evoked release in excitatory and inhibitory synapses without affecting spontaneous release in *unc-31(e928)* in our work hints at a direct involvement of UNC-31 in synaptic vesicle exocytosis. Thus, we examined whether the phenotype of *unc-31* null mutation is occluded by introducing a mutation in EGL-3. EGL-3 is homologue of mammalian type 2 kex2/subtilisin-like proprotein convertase (KPC-2) that is required for processing and subsequent maturation of most *C. elegans* neuropeptides (Kass et al., 2001). Intriguingly, *egl-3(tm1377)* null mutant does not show changes in spontaneous release in both excitatory and inhibitory NMJs, consistent with its main function in peptidergic signaling (**Fig 4A-C, G-I**). Importantly, unlike the *unc-31* mutation, *egl-3(tm1377)* mutation did not decrease evoked release from either excitatory or inhibitory synapses (**Fig 4D-F, J-L**). This suggests that the synaptic transmission defects observed in *unc-31* mutants is at least in part independent of EGL-3-processed peptidergic signaling. Indeed, *unc-31(e928); egl-3(tm1377)* double mutants showed no change in evoked EPSC and IPSC, in comparison to the *unc-31(e928)* single mutants (**Fig 4D-F, J-L**).

**Figure 4.**
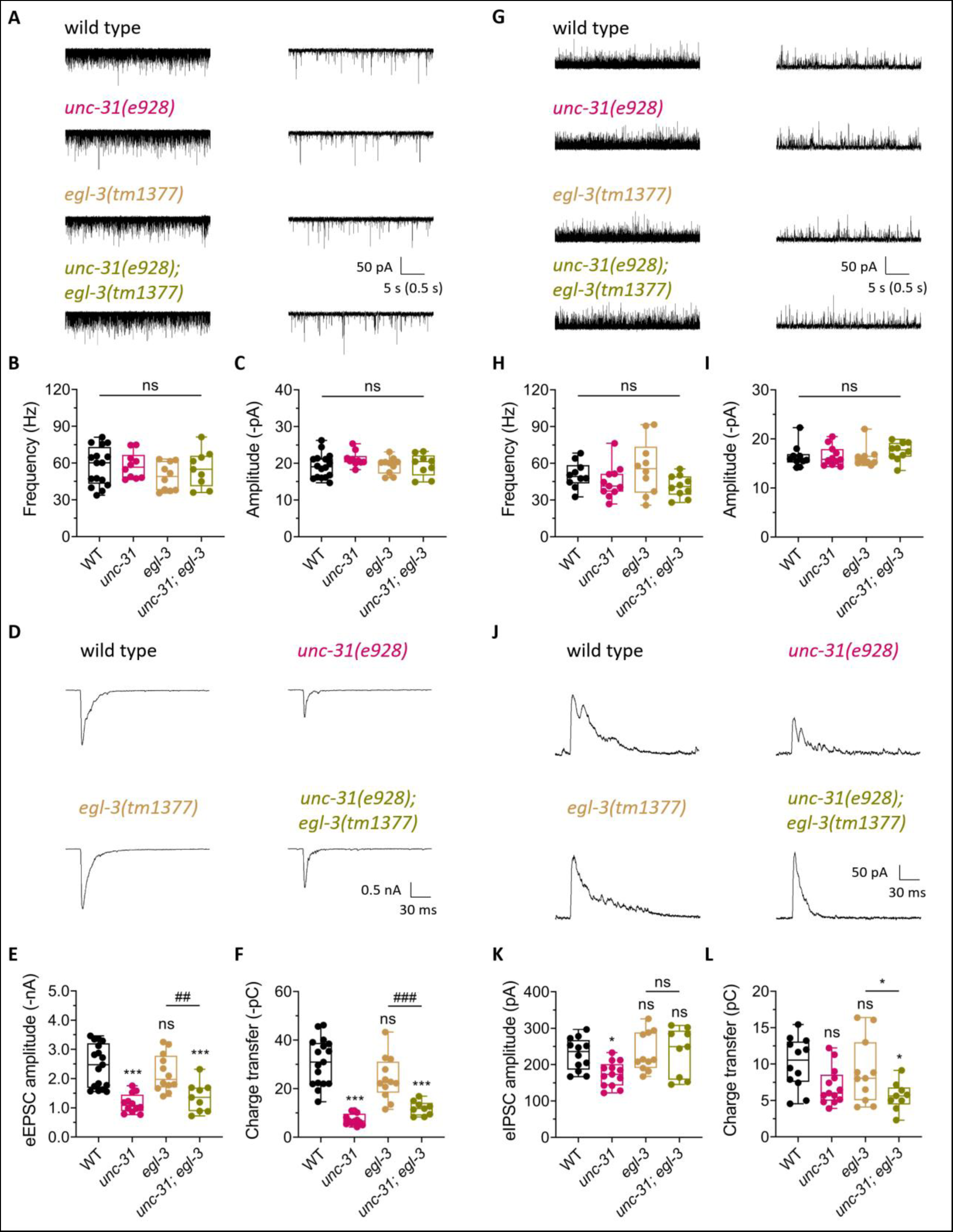
EGL-3-independent regulation of evoked release by UNC-31. (A) Representative traces of mPSCs recorded from WT, *unc-31(e928)*, *egl-3(tm1377),* and *unc-31(e928); egl-3(tm1377)* worms, displayed at different timescales. (B and C) Quantification of the mPSCs frequency (B) and amplitude (C) across different genotypes. (D) Representative traces and quantification of evoked EPSCs recorded from the indicated strains. (G) Representative traces of mIPSCs recorded from the indicated strains, shown at different timescales. (H and I) Quantification of the mIPSCs frequency (H) and amplitude (I) among different genotypes. (J) Representative traces and quantification of evoked IPSCs recorded from the indicated strains. Data are presented as box-and-whisker plots, with the median (central line), 25th–75th percentile (bounds of the box), and 5th–95th percentile (whiskers) indicated. Student’s t-test was performed for comparisons of two groups, ##p<0.01; ###p<0.001; ns, not significant. One-way ANOVA analysis was used for comparisons of multiple groups, followed by Tukey’s range test was performed, *p<0.05; **p<0.01; ***p<0.001; ns, not significant.

We further tested the effects of other enzymes that process peptides, KPC-1, AEX-5, BLI-4, on synaptic vesicle release (**Fig S3**). KPC-1 is homologous to furin proprotein convertase. In addition to its role in processing insulin-like peptides (Hung et al. Development 2014), it was reported to regulate dendritic branching in *C. elegans* (Salzberg et al., 2014; Schroeder et al., 2013). *aex-5* on the other hand encodes a proprotein convertase that regulates the secretion of multiple neuropeptides (Husson et al., 2006; Sheng et al., 2015; Thomas, 1990). Similarly, the *bil-4* gene encodes kex2/substilisin-like endoproteases in the family of proprotein convertase (Thacker et al., 1995). However, none of these proprotein convertase family mutants impair spontaneous or evoked release (**Fig S3**). Therefore, the disruption of neuropeptide synthesis is improbable to detrimentally affect synaptic transmission, thereby suggesting that the modulation of evoked release by UNC-31 is independent and extends beyond its involvement in dense-core vesicle (DCV) regulation.

### *unc-31* null mutation suppresses enhanced spontaneous release in the excitatory synapse of *cpx-1* mutants

After we established evidence on the ability of UNC-31 to directly regulate evoked exocytosis, we set to investigate the relationship between UNC-31 and CPX-1. We generated the double mutant, *cpx-1(ok1552); unc-31(e928)*, and found that they showed a striking decrease in motility, worse than *cpx-1(ok1552)* and *unc-31(e928)* single mutants (**Fig S4A**). Interestingly, *cpx-1(ok1552); unc-31(e928)* demonstrates heightened aldicarb sensitivity, even stronger than *cpx-1(ok1552)* (**Fig S4B**). We hypothesized that the absence of UNC-31 further worsens evoked release and enhances spontaneous release at the excitatory synapses.

We analyzed the excitatory and inhibitory synaptic transmission of the double mutants. As expected, we found that eEPSC and eIPSC are further decreased in double mutant in comparison with each single mutant (**Fig 5D-F, J-L**), suggesting an additive or synthetic effect of *unc-31* and *cpx-1* mutations on evoked release. Similar to the single mutants, *cpx-1(ok1552); unc-31(e928)* decreases evoked release in the excitatory synapses more strongly than in the inhibitory synapse. Therefore, these two proteins appear to be engaged in the regulation of evoked release through distinct pathways in both types of synapses. Unexpectedly, we found that enhanced spontaneous release in the excitatory synapse in *cpx-1(ok1552)* is abolished in *cpx-1(ok1552); unc-31(e928)* double mutant (**Fig 5A-C**). Meanwhile, mIPSC is unaffected in the double mutant (**Fig 5G-I**), consistent with the lack of effects of *unc-31(e928)* and *cpx-1(ok1552)* on mIPSC. Hence, *unc-31* null abolishes the E/I imbalance of *cpx-1* mutant in the spontaneous release. At first, the enhanced sensitivity to aldicarb observed in the double mutant contradicts the pronounced decrease in mPSCs and eEPSCs. The reduction in GABAergic transmission, on the other hand, is able to associate with increased aldicarb sensitivity (Dabbish & Raizen, 2011). Hence, the reduction in inhibition in the evoked release might underlie the hypersensitivity under aldicarb.

**Figure 5.**
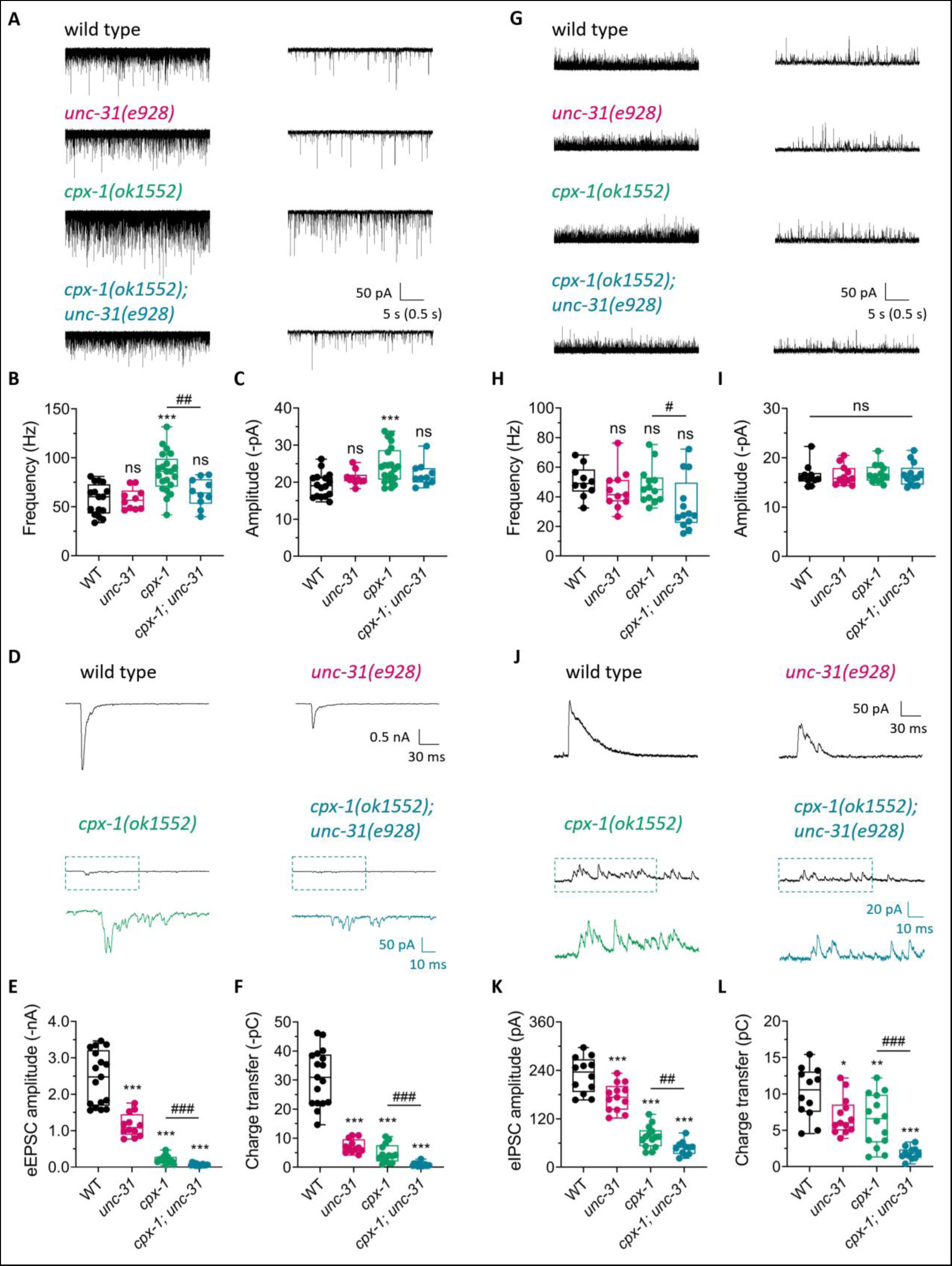
UNC-31 regulates enhanced spontaneous release in cpx-1. (A-C) Representative traces and quantification of mPSCs recorded from the indicated strains. (D-F) Representative traces and quantification of evoked EPSCs recorded from the indicated strains. Zoomed-in regions are displayed in lower traces for *cpx-1(ok1552)* (green) and *cpx-1(ok1552); unc-31(e928)* (blue). (G-I) Representative traces and quantification of mIPSCs recorded from the indicated mutants. (J) Representative traces of evoked IPSCs recorded from the indicated strains. Zoomed-in regions are displayed in lower traces for *cpx-1(ok1552)* (green) and *unc-31(e928); cpx-1(ok1552)* (blue). (K and L) Quantification of the IPSCs amplitude (K) and charge transfer (L) in the same strains. Data are presented as box-and-whisker plots, with the median (central line), 25th– 75th percentile (bounds of the box), and 5th–95th percentile (whiskers) indicated. Student’s t-test was performed for comparisons of two groups, #p<0.05; ##p<0.01; ###p<0.001; ns, not significant. One-way ANOVA analysis was used for comparisons of multiple groups, followed by Tukey’s range test was performed, *p<0.05; **p<0.01; ***p<0.001; ns, not significant. Error bars represent SEM.

We next examined whether *unc-31* null mutation also suppresses enhanced spontaneous release by CPX-1 clamping-specific mutant, *cpx-1(Δ12, syb3584)*. Consistent with our observation from the *cpx-1(ok1552)* null mutants, the enhanced mPSC and more modestly increased mIPSC in *cpx-1(Δ12, syb3584)* mutants are abolished in *cpx-1(Δ12, syb3584); unc-31(e928)* double mutant (**Fig 6A-C, G-I**). *cpx-1(Δ12, syb3584)* clamping mutant further decreases both eEPSC and eIPSC in *unc-31(e928)* null background (**Fig 6D-F, J-L**). Hence, the reduction of exocytosis in *unc-31(e928)* might in part through regulating CPX-1’s clamping function. We conclude that the UNC-31 function is indispensable for maintaining enhanced spontaneous release by the removal of the CPX-1 clamping function (**Fig S5**).

**Figure 6.**
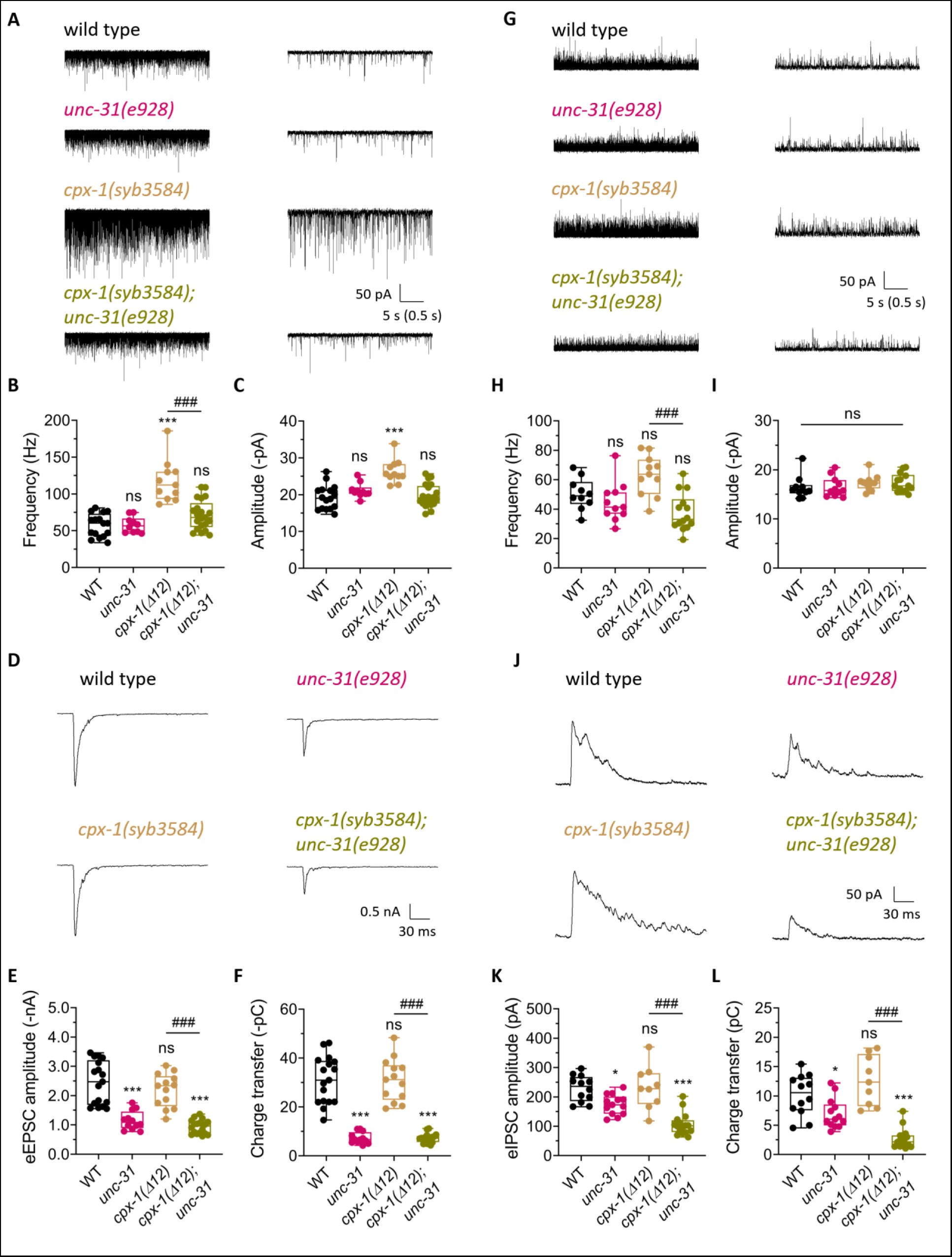
UNC-31 regulates enhanced spontaneous release in cpx-1(Δ12). (A-C) Representative traces and quantification of mPSCs recorded from the indicated genotypes. (D-F) Representative traces and quantification of evoked EPSCs recorded from the indicated genotypes. (G-I) Representative traces and quantification of mIPSCs recorded from the indicated genotypes. (J-L) Representative traces and quantification of evoked IPSCs recorded from the indicated genotypes. Data are presented as box-and-whisker plots, with the median (central line), 25th–75th percentile (bounds of the box), and 5th–95th percentile (whiskers) indicated. Student’s t-test was performed for comparisons of two groups, ###p<0.001; ns, not significant. One-way ANOVA analysis was used for comparisons of multiple groups, followed by Tukey’s range test was performed, *p<0.05; ***p<0.001; ns, not significant. Error bars represent SEM.

### The suppression of *cpx-1* by *unc-31* relies on the presence of external Ca^2+^

Spontaneous release of SVs requires external Ca^2+^. We then examined the external Ca^2+^-dependence of the enhanced mPSC in *cpx-1* mutant (**Fig S6**). At 5 mM external Ca^2+^, the spontaneous release frequency in *cpx-1* mutant is ~2-fold of that in wild type animals (**Fig S6A, B**). In both *cpx-1* mutants and wild-type animals, removal of external Ca^2+^ reduced the mPSC frequency, whereas a 30 min pre-incubation with 1 mM BAPTA-AM in Ca^2+^-free medium lowered the mPSC frequency >4-fold (**Fig S6A, B**). The increased mPSC amplitude was not reduced in the absence of external Ca^2+^ with or without pre-incubation of BAPTA-AM (**Fig S6C**). Thus, consistent with prior studies (Yang et al., 2011), the enhanced mPSCs observed in *cpx-1* mutant are Ca^2+^-dependent.

UNC-31 encodes a “Calcium-dependent Activator Protein for Secretion, CAPS” protein. The dependency of CAPS/UNC-31 function on Ca^2+^ for its role in regulating synaptic vesicle exocytosis has been extensively studied. In our study, we have discovered that UNC-31 plays a crucial role in maintaining the enhanced spontaneous release observed in *cpx-1* mutants. This finding prompts the question of whether UNC-31 functions as the secondary Ca^2+^-sensor in addition to synaptotagmin-1, as previously hypothesized from complexin knock-down experiments (Yang et al., 2010). To explore this hypothesis, we investigated the Ca^2+^ dependency of UNC-31’s regulatory effect on *cpx-1* mutants. At external Ca^2+^ from 0.5 mM to 5 mM, a comparable mPSC pattern to wild-type was evident in the *unc-31* single mutant. In contrast, the increased mPSC frequency and amplitude observed in *cpx-1* mutants were abolished in *cpx-1; unc-31* double mutants, providing the confirmation of *unc-31*’s inhibitory influence on *cpx-1*. Interestingly, under conditions of zero external Ca^2+^, the enhanced mPSCs in *cpx-1* displayed no significant changes in either frequency or amplitude when compare to *cpx-1; unc-31* double mutants. (**Fig S6D-F**). These results implies that the suppression of *cpx-1* by *unc-31* exhibits an obvious dependency on the presence of external Ca^2+^. Alternatively, CAPS/UNC-31 could potentially function as the Ca^2+^-sensor responsible for spontaneous release, a role that is hindered by the presence of complexin/CPX-1.

### Stimulus-dependent persistent increase of spontaneous release is diminished in *unc-31* mutant

To investigate the functions of UNC-31 in sustaining enhanced spontaneous mPSC physiologically, we tested whether activity-dependent synaptic enhancement is affected in *unc-31(e928)* mutant. We applied 10 trains of 0.1s light stimulation to the excitatory synapses (**Fig 7A**). 10s after the modest stimuli train, the wild type showed an increase in mPSC frequency (**Fig 7A, C**). Nevertheless, the stimulation train does not enhance spontaneous release in the *unc-31(e928)* (**Fig 7A, C**). Moreover, we applied a tonic stimulation (10 trains of 1s stimulation) and found that mPSC frequency is further enhanced in the wild type but not in the *unc-31(e928)* (**Fig 7B, D**). We tested the short-term depression of the evoked EPSC (normalized to the first eEPSC) after modest and tonic stimulation and found no difference between wild type and *unc-31(e928)* (**Fig S7**). Thus, UNC-31 becomes critical to maintain spontaneous release when it is enhanced by both genetic mutation and repetitive stimulation.

**Figure 7.**
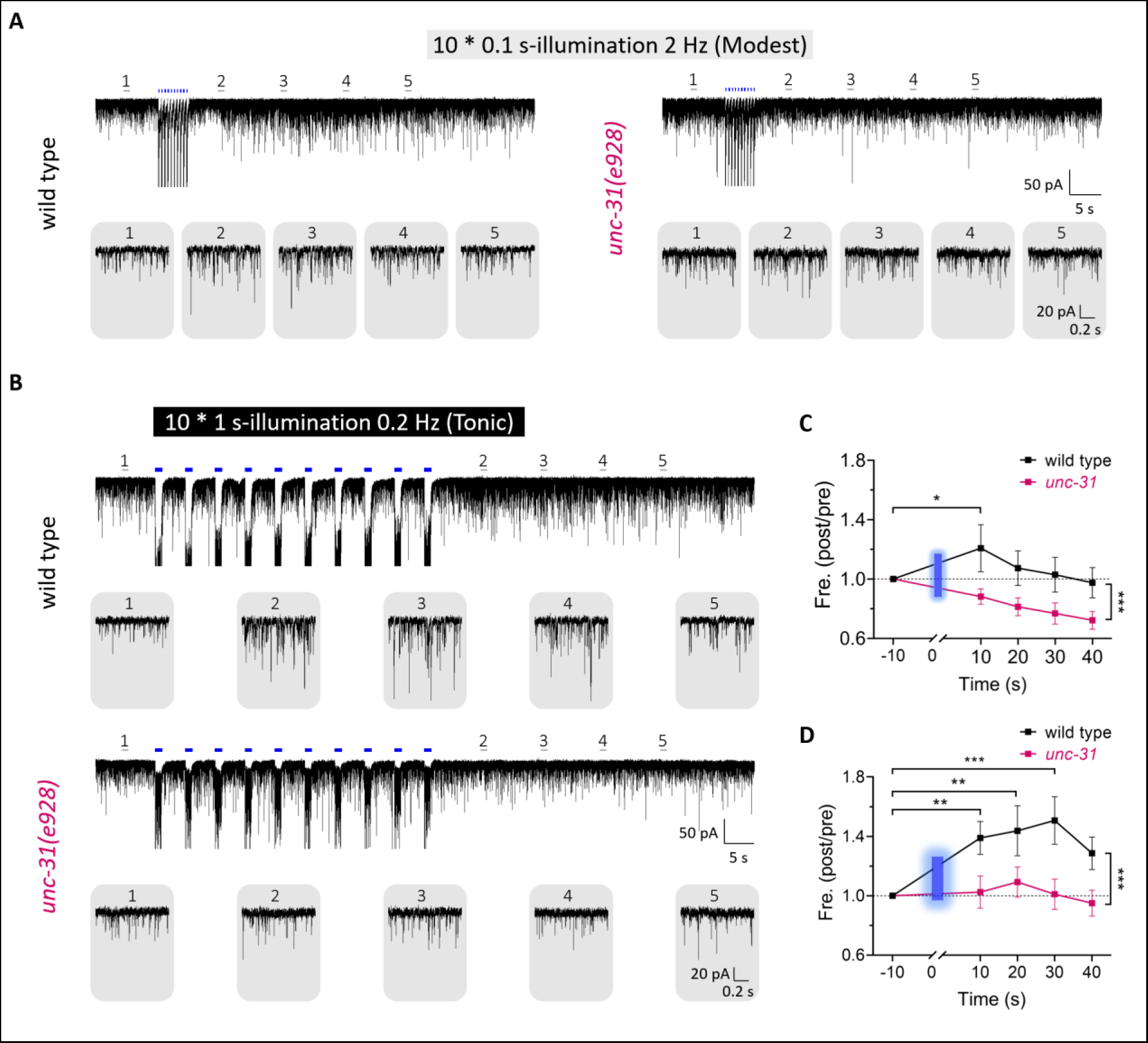
UNC-31 requirement for stimuli-dependent spontaneous release increase. (A) Sample traces of persistent mPSCs evoked by repeated blue light-stimulation (blue bars: 10 x 0.1 s pulses at 2 Hz) from WT and *unc-31(e928)* worms. Zoomed-in views of individual minis traces are presented in the gray rounded rectangle, extracted from indicated regions of the upper panel (lines 1-5). (B) Sample traces of mPSCs before and after the stimulation by repeated stimulation (blue bars: 10 x 1 s pulses at 0.2 Hz). Subsequent persistent enhanced spontaneous releases were recorded from WT and *unc-31(e928)* worms. (C) Ratio between mPSCs frequency post- and pre-blue light illumination of (A). The blue bars depict the duration of 10 consecutive illumination. (D) Ratio between mPSCs frequency post- and pre-blue light illumination of. The blue bars represent 10 consecutive illumination (B). Two-way ANOVA analysis was performed, *p<0.05; **p<0.01; ***p<0.001. Error bars represent SEM.

## Discussion

Much of our understanding of regulating SNARE-dependent synaptic vesicle release comes from excitatory synapses, in part due to technical difficulties, while assuming their functions are preserved in inhibitory neurons. Our results demonstrated the unexpected differential regulation of excitatory and inhibitory synaptic release by CPX-1 and UNC-31. Most strikingly, the increase in spontaneous release in the *cpx-1* null mutant is selective in the excitatory synapses. We also found that *the unc-31 null* mutant, independent of its DCV regulation, impairs evoked release more severely in the excitatory synapses than the inhibitory synapses. Thus, we warren further investigation of the mechanism of exocytosis in inhibitory synaptic releases. The differential regulation of exocytosis will help understand the molecular mechanisms underlying disrupted E/I balance in diseases and impaired brain functions.

Complexin can facilitate evoked release and clamps spontaneous asynchronous release (Iyer et al., 2013; Maximov et al., 2009). Here, we show that CPX-1 functions depend on the synapses. In the *cpx-1* null mutant, evoked release is decreased in both excitatory and inhibitory synapses, whereas spontaneous release is increased only in the excitatory synapses. Our finding is consistent with the previous literature on complexin in mice and *Drosophila* showing the most consistent phenotype of complexin mutants is reduced evoked release rather than enhanced spontaneous release in excitatory transmission (Iyer et al., 2013; Jorquera et al., 2012; Xue et al., 2008). Regarding GABAergic transmission, complexin knockout in mammalian striatal GABAergic neuronal culture shows a decrease in evoked and spontaneous IPSC (Xue et al., 2008). While our results are consistent with the decreased evoked release, but inconsistent with the effects observed in spontaneous release. The inconsistencies come partly from differences across species, where invertebrates CTD has stronger inhibitory effects compared to mammalian complexin (Wragg et al., 2017; Xue et al., 2009). To this end, species variation on the functions of complexin in controlling inhibitory synaptic vesicles released needs to be clarified. As locomotion depends on the E/I balance at the NMJ (Thapliyal & Babu, 2018), the tilted balance towards increased inhibition explains the impaired movement in the *cpx-1* mutant. Reduction in complexin level is found in Schizophrenia, major depressive disorder, and neurodegenerative diseases such as Huntington’s and Alzheimer’s diseases (Brose, 2008). Hence, we propose that the loss of complexin alters E/I balance in the neural circuits as a mechanism of brain disorders.

The clamping function of complexin/CPX-1 has been considered to be critical to prevent the depletion of primed vesicles that allow synchronized evoked release (Chang et al., 2015; Vaithianathan et al., 2015). However, our finding of the clamping-specific mutant, *cpx-1(Δ12)*, shows normal evoked release despite the dramatically enhanced spontaneous release in the excitatory synapse and a modest increase in the inhibitory synapse (**Fig 2**). A similar clamping-specific mutant has been reported in *Drosophila*, yet the specificity on regulating spontaneous effects is spliced isoform dependent (Buhl et al., 2013; Iyer et al., 2013). Therefore, clamping of spontaneous release is not a prerequisite to support evoked release. Works in both mammalian and *Drosophila* excitatory and inhibitory synapses suggest spontaneous and evoked release can be segregated, with differential pools of synaptic vesicles for each mode of release at the active zone (Horvath et al., 2020; Melom et al., 2013). Hence, CPX-1 might as well participate in the regulation of vesicle release in both pools. Previous biochemical work showed that synaptotagmin is the one that removes the complexin clamping to trigger synchronized release (Bera et al., 2022; Ramakrishnan et al., 2020; Zhou et al., 2017). To our surprise, genetic evidence to support this well-known hypothesis is missing and contradictory. A double mutant in *Drosophila* with the loss of both complexin and synaptotagmin shows an additive effect in impairing spontaneous and evoked release, indicating partly independent regulatory functions (Jorquera et al., 2012). However, in mice, the synaptotagmin Ca^2+^-binding C2B domain can clamp synaptic vesicles independent of complexin (Courtney et al., 2019). More work is necessary to clarify the genetic and functional interaction between synaptotagmin and complexin.

UNC-31 is well established in its role in DCV regulation of neuropeptide release but controversial in synaptic vesicle regulation (Cai et al., 2004; Charlie et al., 2006; Lin et al., 2010; Speese et al., 2007; Zhou et al., 2007). We found that *unc-31* null mutation decreased evoked exocytosis without affecting spontaneous release, and this decrease was more evident in the excitatory synapse. Previously, the disrupted synaptic transmission of *unc-31* mutants is considered secondary to its defects of DCV secretion (Gracheva et al., 2007). However, we found that none of the mutants that play important roles in DCV maturation or release mimicked or occluded the phenotype of *unc-31*. We also presented direct evidence of UNC-31 regulating GABAergic synaptic vesicle clustering. This is consistent with mammalian CAPS function in docking synaptic vesicles (Imig et al., 2014; Jockusch et al., 2007). Thus, contrary to the previous notions in *C. elegans*, functions of UNC-31 should be revisited in direct regulation of synaptic vesicle release. *Unc-31* mutant is immobile and appears in a relaxed posture (Avery et al., 1993). Based on our findings we propose the tilted balance towards stronger inhibition drives the impaired motor behavior.

Interestingly, although *unc-31* null mutation does not affect spontaneous release, it blocks the enhanced spontaneous release caused by *cpx-1* mutants, as well as under repetitive depolarization stimuli. Thus, UNC-31 might play a critical role in short-term plasticity and synaptic depression. CAPS-1 deletion in the thalamocortical synapse causes earlier synaptic depression under repetitive stimuli (Nestvogel et al., 2020). This is consistent with our observation that under tonic stimulation, *unc-31* null is unable to increase spontaneous synaptic vesicle release. Thus, it is tempting to postulate that synaptic vesicle clustering or priming by UNC-31 is not only important for evoked synaptic release but also for the maintenance of enhanced spontaneous release under synaptic plasticity and in synapses with a high release probability.

By studying the genetic interaction between *unc-31* and *cpx-1*, we also observed interesting interplays between the two regulatory proteins in synaptic functions. While the double mutant decreases evoked release strongly without affecting the spontaneous release in excitatory and inhibitory transmission, it has a very high aldicarb sensitivity. Aldicarb as an acetylcholinesterase inhibitor, induces paralysis in nematodes with high acetylcholine release. However, the reduction of GABAergic transmission also increases aldicarb sensitivity (Dabbish & Raizen, 2011; Locke et al., 2009; Vashlishan et al., 2008). Therefore, we can interpret the aldicarb hypersensitivity in E/I imbalance, with both evoked EPSC and IPSC being strongly abolished. In imaging the synaptic vesicle clustering, we found that the deletion of *cpx-1* in the *unc-31* background rescued *the unc-31* phenotype of disrupted vesicular docking. Given the additive effects of decreased evoked release, it is unclear how vesicular clustering is rescued. As UNC-31 involves primarily in docking while CPX-1 plays a role in priming after the docking step (Imig et al., 2014; Zhou et al., 2019), the increase in synaptic vesicle clustering may indirectly suggest impairment in priming leading to vesicular accumulation. The abolished evoked release may lead to an increase in unfused vesicles at the terminal, which may lead to the recovery or rescue of reduced vesicle clustering observed in the *unc-31* single mutant. Therefore, UNC-31 and CPX-1 work independently in regulating the docking and priming steps respectively in vesicular fusion.

Together, our results using optogenetics revealed unexpected E/I imbalances in previously studied synaptic exocytosis mutants and our new clamping-specific KI mutant. We demonstrated the importance of studying inhibitory synaptic release whose mechanism may be significantly different from that in excitatory synapse. The precise electrophysiological analysis of synaptic transmission mutants in both excitatory and inhibitory synapses will be critical for a better understanding of the behavior of the respective mutants. With increasing knowledge in the regulation of excitatory and inhibitory synapses, we will gain better access to targets manipulating E/I balance in neural circuits underlying neurological and psychiatric diseases.

## Materials and Methods

### Strains

The complete lists of constructs, transgenic lines and strains generated or acquired for this study are provided in the Supplementary Table 1. All *C. elegans* strains were cultured on standard Nematode Growth Medium (NGM) plates seeded with OP50, and maintained at 22 °C. Unless stated otherwise noted, the wild-type animal refers to the Bristol N2 strain. Only hermaphrodite worms were used for the experiments.

### Aldicarb assays

Aldicarb sensitivity was assessed using synchronously grown adult worms placed on non-seeded 30 mm NGM plates containing 0.3 mM or 1 mM aldicarb. All assays were done in 1 mM aldicarb plates unless specified otherwise. Over a 4 or 24-hour period, worms were monitored for paralysis at 15- or 30-min intervals. Worms were considered paralyzed when there was no movement or pharyngeal pumping in response to 3 taps to the head and tail with a platinum wire. Once paralyzed, worms were removed from the plate. 6 sets of 15-20 worms were examined for each strain.

### Thrashing assay

Motility of each strain was determined by counting the thrashing rate of *C. elegans* in liquid M9 medium. Briefly, worms were bleached to release their eggs and were then synchronously grown to young adulthood. Young adult worms were placed in a 60 μL drop of M9 buffer on a 30 mm petri dish cover. After a 2-minute recovery period, worms were video-recorded for 2 minutes using an OMAX A3580U camera on a dissecting microscope with the OMAX ToupView software. The number of thrashes per minute were manually counted and averaged within each strain. A thrash was defined as a complete bend in the opposite direction at the midpoint of the body.

### Fluorescence Microscopy

Images of fluorescently tagged proteins were captured in live L4 worms. L4 stage transgenic animals expressing different fluorescence markers were prepared a day before imaging. Worms were immobilized by 2.5 mM levamisole (Sigma-Aldrich, USA) in M9 buffer solution. Fluorescence signals were captured using a Plan-Apochromatic 60X objective on a confocal microscope (FV3000, Olympus, Japan) in the same conditions. Experiments were performed at room temperatures (20-22 °C).

### In-situ electrophysiology

Dissection and recording were carried out using protocols and solutions described in the previous studies (Gao & Zhen, 2011; Richmond & Jorgensen, 1999). Briefly, 1- or 2-day-old hermaphrodite adults were glued (Histoacryl Blue, Braun) to a sylgard (DowCorning, USA)-coated cover glass covered with bath solution. Under a DIC microscope, semi - fixed worms were dissected dorsally with a glass pipette, the cuticle flap was flipped and gently glued (WORMGLU, GluStitch Inc.) to the opposite side, and the intact ventral body muscles were exposed after cleaning the viscera. Anterior body wall muscle cells were patched using 4-6 MΩ resistant borosilicate pipettes (World Precision Instruments, USA), which were pulled by micropipette puller P-1000 (Sutter). Membrane currents were recorded in the whole-cell configuration by PULSE software with an EPC-9 patch clamp amplifier (HEKA, Germany). Data were digitized at 10 kHz and filtered at 2.6 kHz. The pipette solution contains (in mM): K-gluconate 115; KCl 25; CaCl_2_ 0.1; MgCl_2_ 5; BAPTA 1; HEPES 10; Na_2_ATP 5; Na_2_GTP 0.5; cAMP 0.5; cGMP 0.5, pH 7.2 with KOH, ~320 mOsm. cAMP and cGMP were included to maintain the activity and longevity of the preparation. The bath solution consists of (in mM): NaCl 150; KCl 5; CaCl_2_ 5; MgCl_2_ 1; glucose 10; sucrose 5; HEPES 15, pH 7.3 with NaOH, ~330 mOsm. Different extracellular Ca^2+^ bath solutions were prepared according to the bath solution with different CaCl_2_ (0, 0.5, 1.5, 5 mM). Zero Ca^2+^ with BAPTA solution consists of additional 1 mM BAPTA in the 0 Ca^2+^ bath solution to further chelate and reduce the extracellular free [Ca^2+^] (Yang et al., 2010). Muscle cells were held at −60 mV when recording mPSCs or eEPSCs. To isolate mIPSCs or eIPSCs, recordings were performed with a holding potential of −10 mV, with 0.5 mM D-tubocurarine (d-TBC) included in the bath solution to block all acetylcholine receptors (Richmond & Jorgensen, 1999; Maro et al., 2015). Chemicals were obtained from Sigma unless stated otherwise. Experiments were performed at room temperatures (20-22 °C).

### Synaptic vesicles imaging

Worms were bleached to release their eggs, L1-stage were picked ~8-9 hours after and were grown to L4-stage. Live L4 worms were mounted on 2% dry agarose pads with 1μL of M9 buffer and imaged with a spinning disk confocal microscope at 63x oil objective in the region between the RVG and the vulva. For each worm, Z stacks were taken of the nerve cord. Images were analyzed using a punctaanalyser toolkit developed in-house (described in (Hung et al., 2007; Kim et al., 2008)) and a custom MATLAB program was used to analyze the images.

### Statistical analysis

The two-tailed Student’s *t*-test was used to compare two groups of data sets. One way or two way ANOVA analysis was used to compare the significant different more than two groups. *P* < 0.05 was considered to be statistically significant; *, ** and *** denote *P* < 0.05, *P* < 0.01, *P* < 0.001, respectively. Graphing and subsequent analysis were performed using Igor Pro (WaveMetrics), Clampfit (Molecular Devices), Image J (National Institutes of Health), Python, Matlab (MathWorks, USA), GraphPad Prism 8 (GraphPad Software Inc., USA) and Excel (Microsoft, USA). For behavior analysis, calcium imaging, electrophysiology and fluorescence imaging, each recording trace was obtained from a different animal. Unless specified otherwise, data were presented as the mean ±SEM.

## Author Contributions

S.G. and S.S. conceived experiments and wrote the manuscript. Y.W., C.C., M.H., performed experiments and analyzed data. R.H., N.R., and C.Z. contributed to the experiments.

## Acknowledgements

We thank Mei Zhen for reagents, strains, valuable inputs and comments, Josep Rizo for insightful comments and discussion, Xia-Jing Tong for strains and helpful suggestions, Lili Chen and Xuhui Chen for experimental supports and data analysis. This research was supported by the Major International (Regional) Joint Research Project (32020103007 to S.G.), the National Natural Science Foundation of China (32371189, 31871069 to S.G.), the National Key Research and Development Program of China (2022YFA1206001 to S.G.), Natural Sciences and Engineering Research Council of Canada (RGPIN 2020 07139 to S.S.) and the Canadian Institute of Health Research (CIHR PJT 165917 to S.S.), the Overseas High-level Talents Introduction Program. We thank *Caenorhabditis Genetics Center*, which is funded by the NIH Office of Research Infrastructure Programs (P40 OD010440), for strains.

---

**Table 1.**
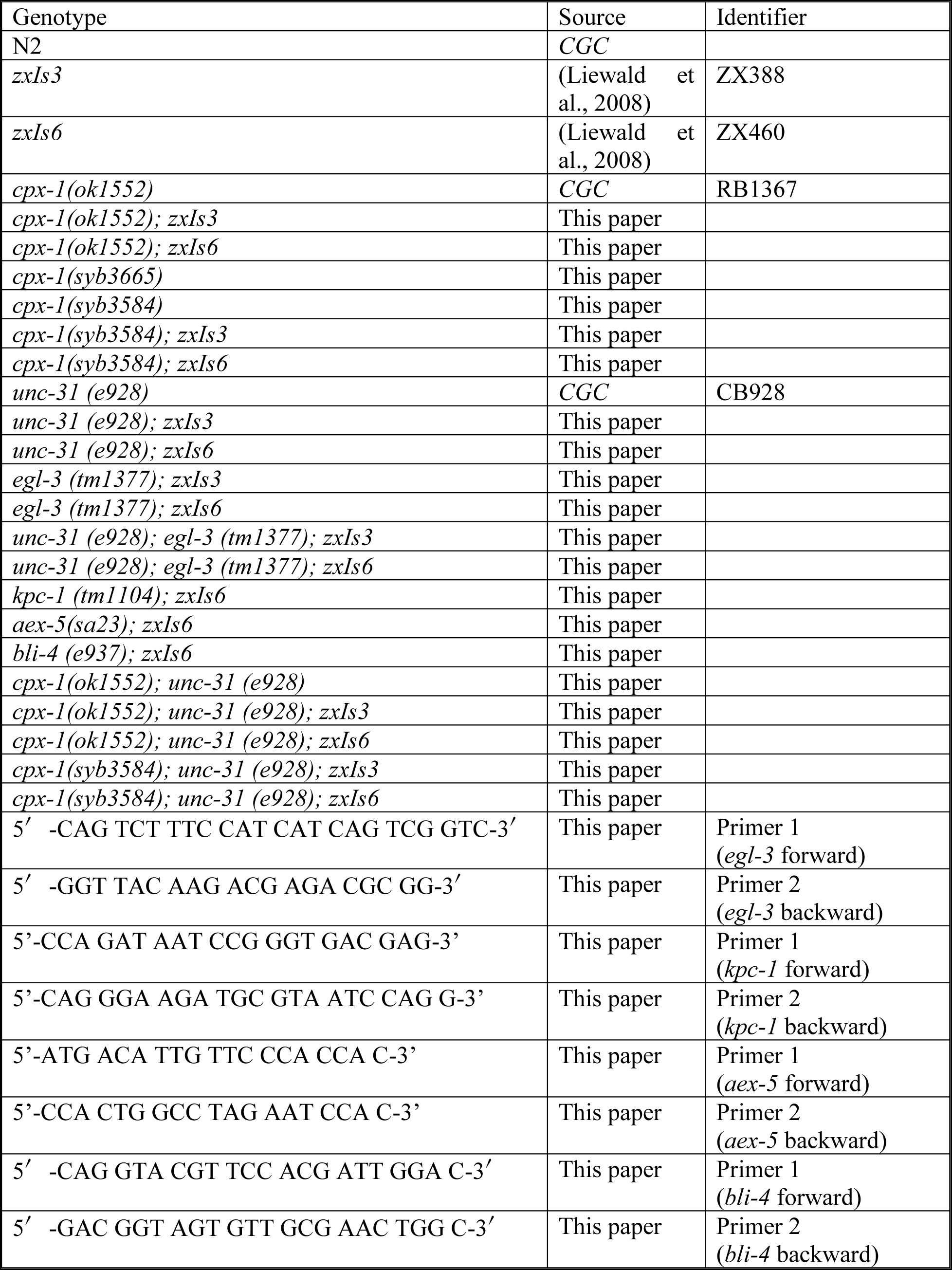

**Supplementary Figure S1.**
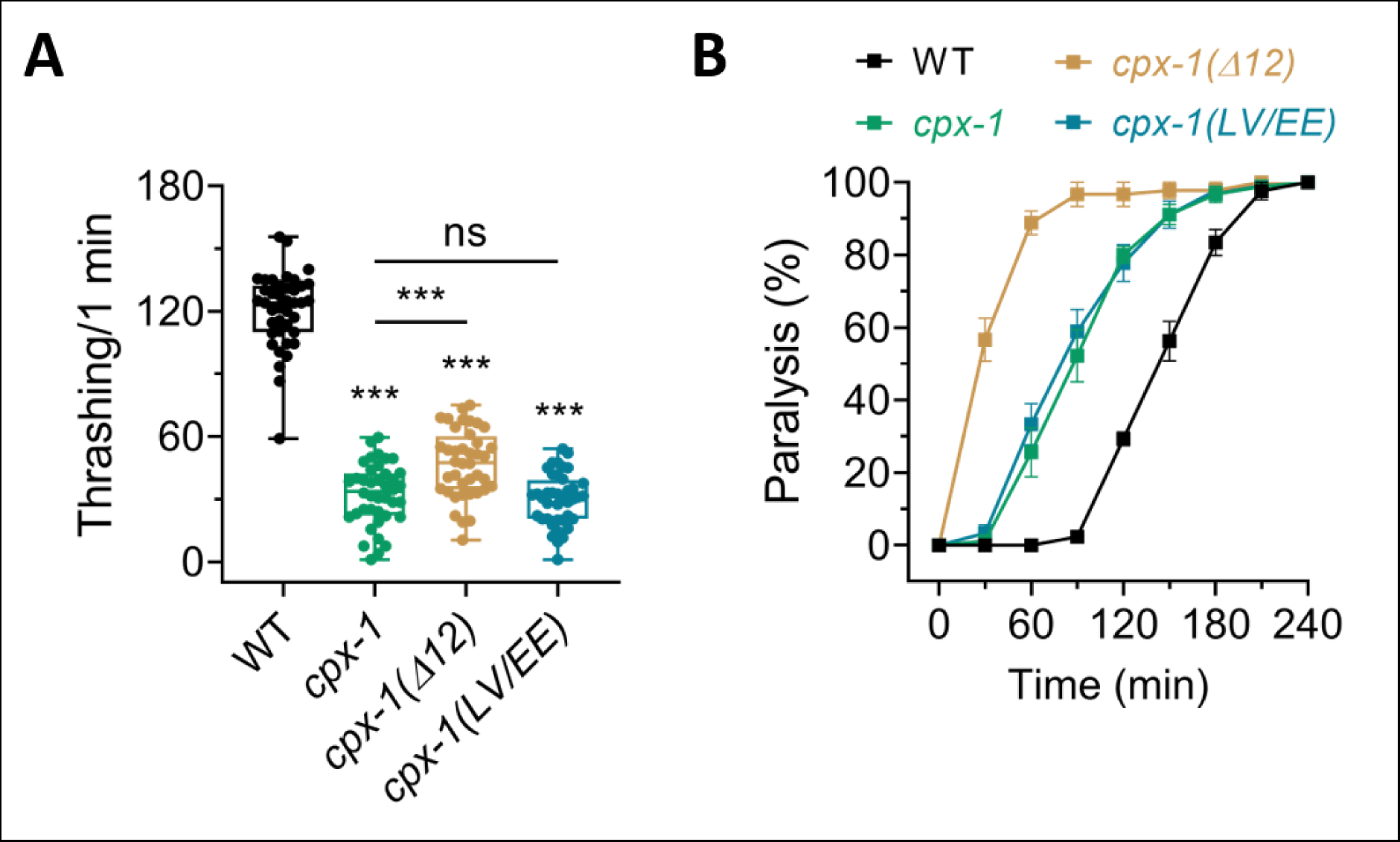
Impaired motility and aldicarb hypersensitivity in cpx-1 mutants. (A) Motility measured by thrashing number per minute across wild type, *cpx-1(ok1552)*, *cpx-1(Δ12, syb3584)*, and *cpx-1(LV/EE, syb3665)*. The analysis included 40 individuals from each strain. (B) Aldicarb assay for indicated strains. Six trials were conducted with 15 worms in each trial. One-way ANOVA analysis was used for comparisons of multiple groups, followed by Tukey’s range test was performed, ***p<0.001; ns, not significant. Error bars represent SEM.

**Supplementary Figure S2.**
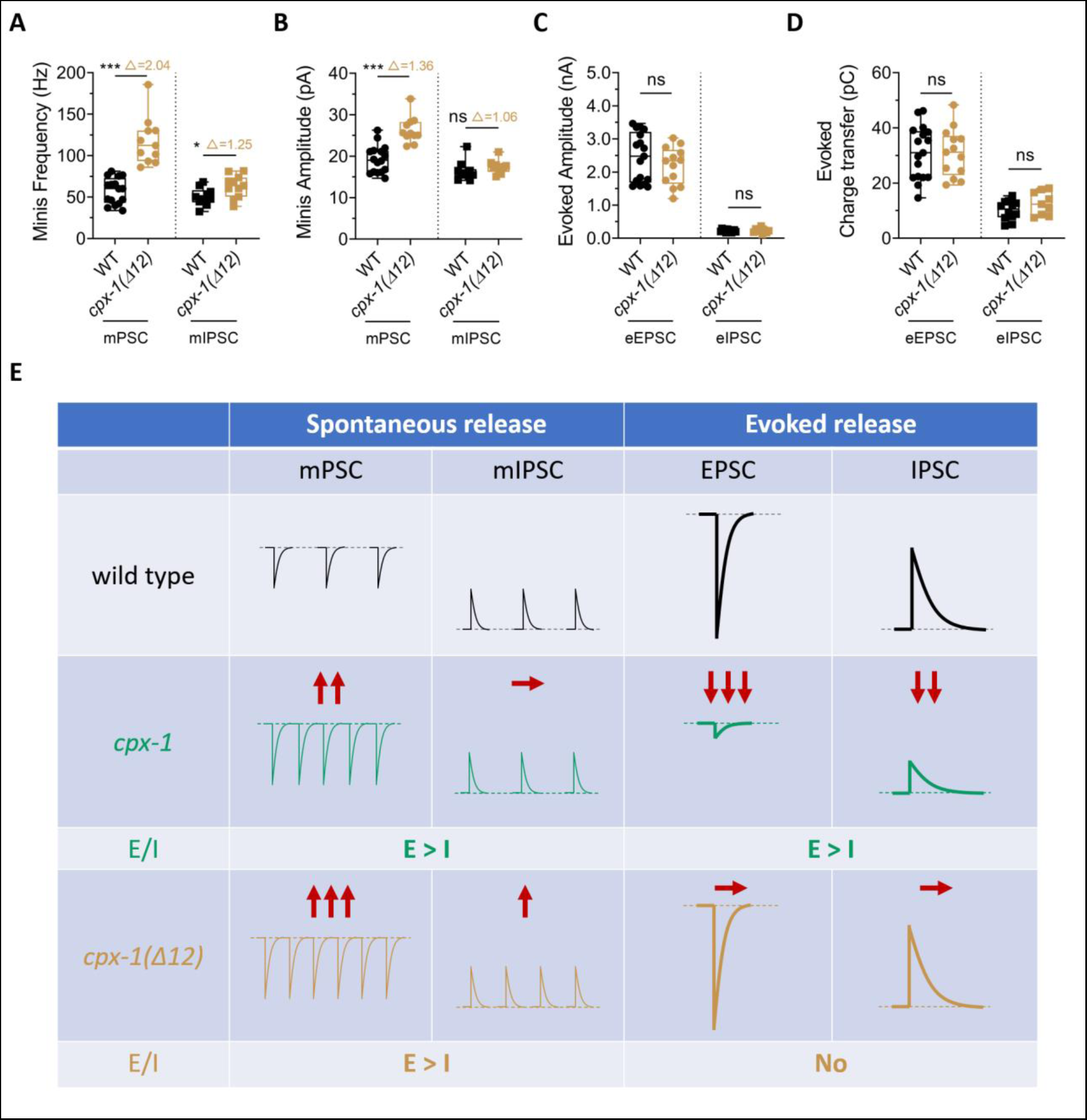
Enhancement of excitatory spontaneous release by cpx-1(Δ12) mutation without impact on evoked release. (A and B) Box-and-whisker plots of the mPSCs (left)/mIPSCs (right) frequency (A) and amplitude (B) in WT and *cpx-1(Δ12)* worms. The numbers after Δ represent the ratio of the average values between WT and *cpx-1(Δ12)*. (C and D) Box-and-whisker plots of the EPSCs (left)/IPSCs (right) amplitude (C) and charge transfer (D) in WT and *cpx-1(Δ12)* worms. (E) Graph depicting the differential regulation of excitatory and inhibitory synaptic vesicle release by the two *cpx-1* mutants. Data are presented as box-and-whisker plots, with the median (central line), 25th–75th percentile (bounds of the box), and 5th–95th percentile (whiskers) indicated. Student’s t-test was performed, *p<0.05; ***p<0.001; ns, not significant.

**Supplementary Figure 3.**
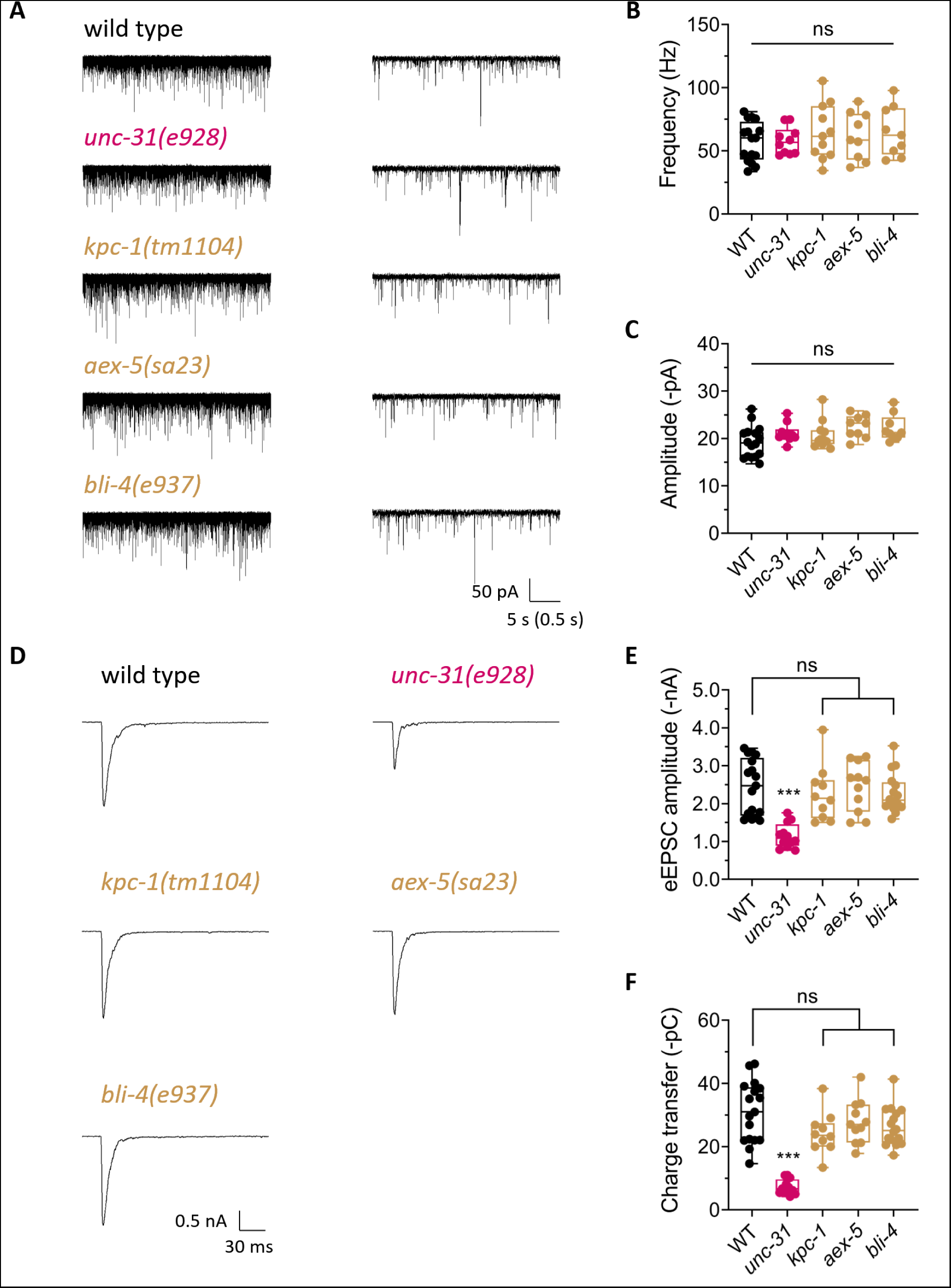
Evoked excitatory neurotransmission regulation of UNC-31 is independent on GPCR downstream signaling pathways. (A) Representative mPSCs traces recorded from WT, *unc-31(e928)*, *kpc-1(tm1104)* and *aex-5(sa23); bli-4(e937)* worms. Right panels show a 0.5 s scale bar for clarity. (B and C) Quantification of the mPSCs frequency (B) and amplitude (C) across the indicated genotypes, as shown in panel (A). (D-F) Representative traces and quantification of evoked EPSCs recorded from the mentioned genotypes. Data are presented as box-and-whisker plots, with the median (central line), 25th–75th percentile (bounds of the box), and 5th–95th percentile (whiskers) indicated. One-way ANOVA analysis was performed, ***p<0.001; ns, not significant.

**Supplementary Figure S4.**
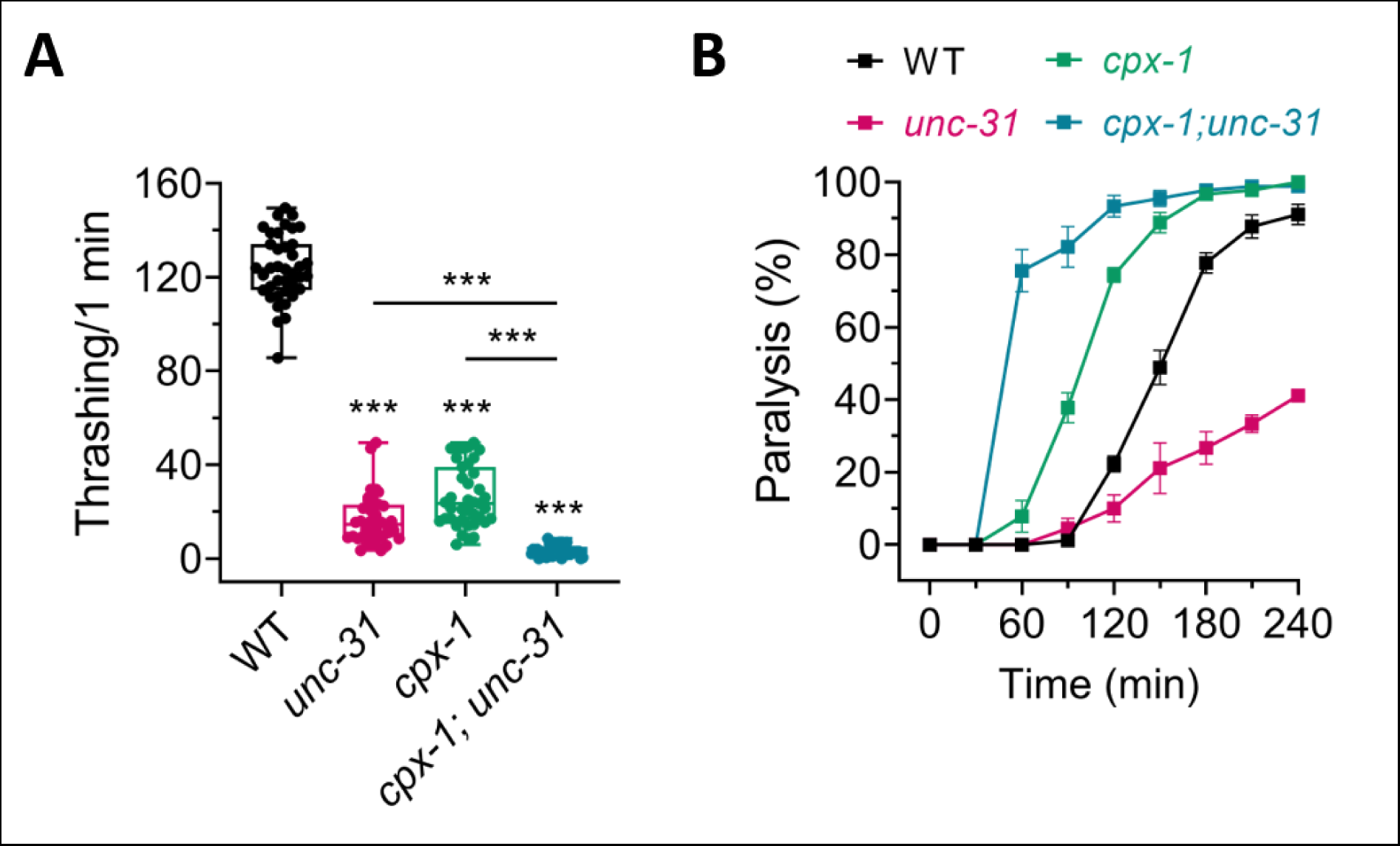
Exacerbated motor impairment and aldicarb hypersensitivity in cpx-1(ok1552); unc-31(e928) double mutants. (A) Motility measured by thrashing number per minute. n = 40 in each strain. (B) Aldicarb assay for wild type, *cpx-1(ok1552)*, *unc-31(e928)*, and the double mutant. n = 6 trials with 15 worms in each trial. Data are mean ±SEM. One-way ANOVA analysis was performed. ***p<0.001.

**Supplementary Figure S5.**
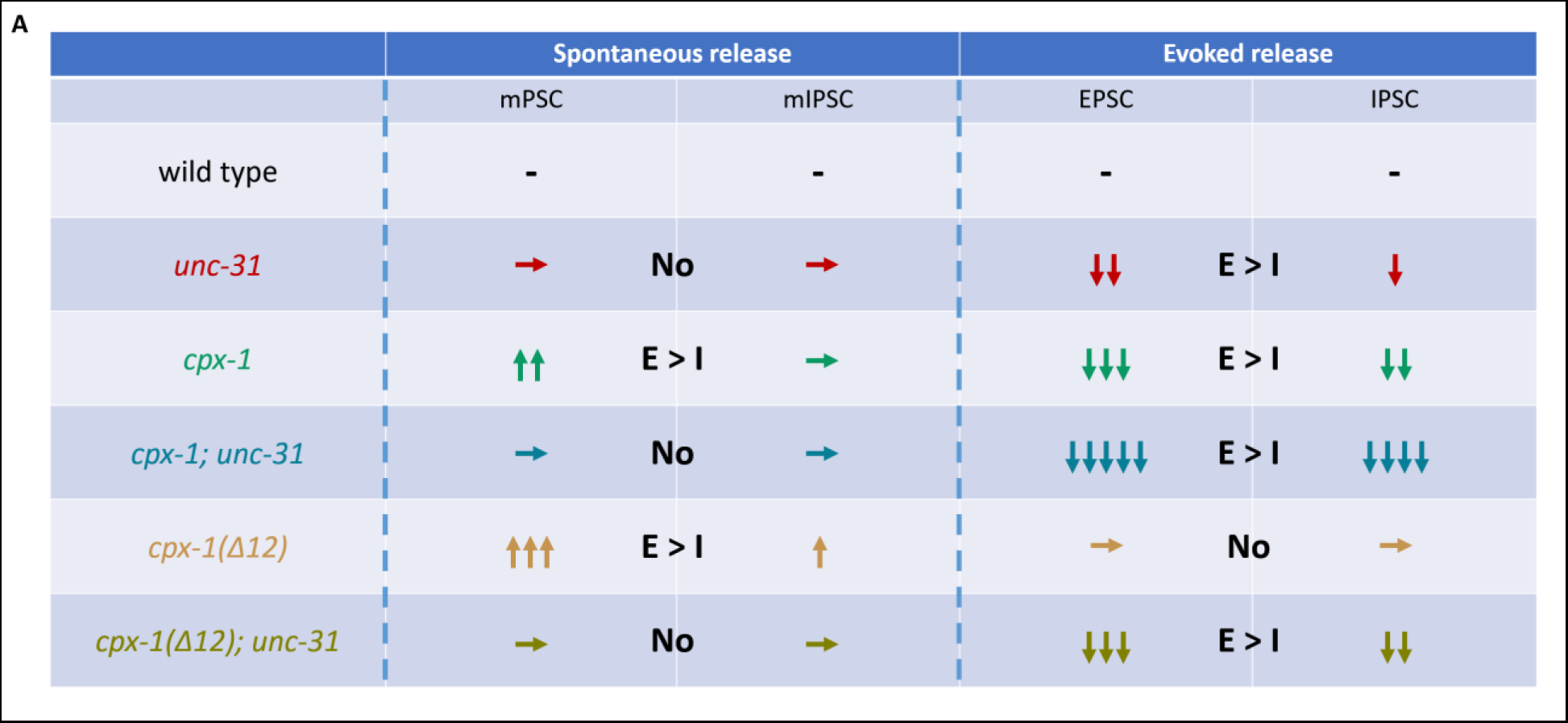
UNC-31 regulates enhanced spontaneous release in cpx-1 mutants. (A) Summary graph showing the differential regulation of excitatory and inhibitory synaptic vesicle release of the indicated genotypes.

**Supplementary Figure S6.**
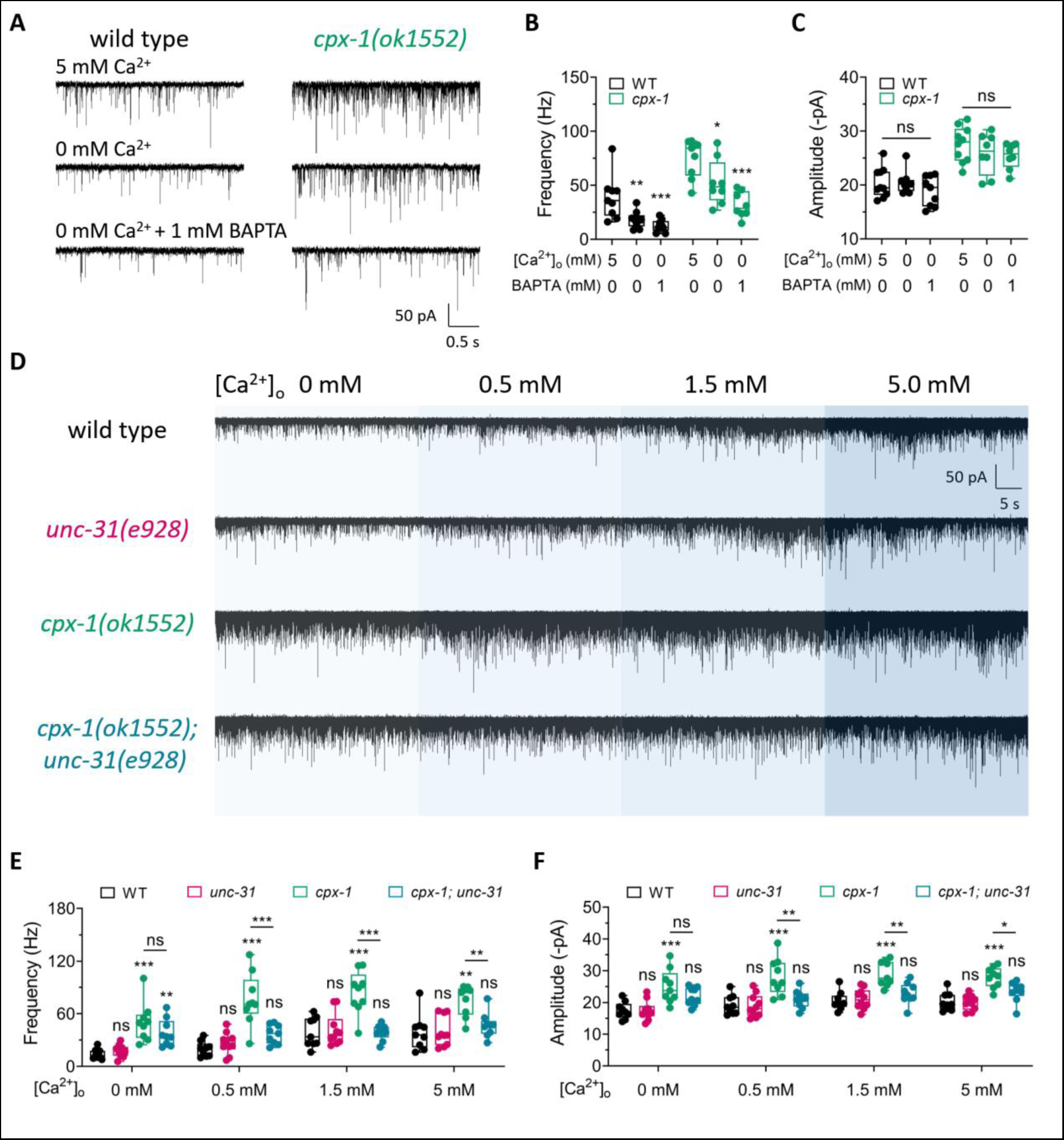
Ca^2+^-dependent suppression of the enhanced mPSCs in cpx-1 mutants by UNC-31. (A) Representative traces of mPSCs recorded from WT and *cpx-1(ok1553)* worms in bath solution containing 5 mM [Ca^2+^]_o_, 0 mM [Ca^2+^]_o_ and 0 mM [Ca^2+^]_o_ with 1 mM BAPTA, respectively. (B, C) Quantification of mPSCs frequency and amplitude of WT and *cpx-1(ok1553)* in various bath solutions. The evident Ca^2+^ dependence of enhanced mPSCs in *cpx-1* mutants is observed. (D) Representative traces of mPSCs recorded from WT, *unc-31(e928)*, *cpx-1(ok1553)* and *cpx-1(ok1553); unc-31(e928)* worms in different Ca^2+^ bath solutions. (E, F) Quantification of mPSCs frequency and amplitude of the indicated strains. The suppression of the enhanced spontaneous release of *cpx-1* by *unc-31* was only apparent in the presence of [Ca^2+^]_o_. Two-way ANOVA analysis was performed. ns, no significance, *p<0.05, **p<0.01, ***p<0.001.

**Supplementary Figure S7.**
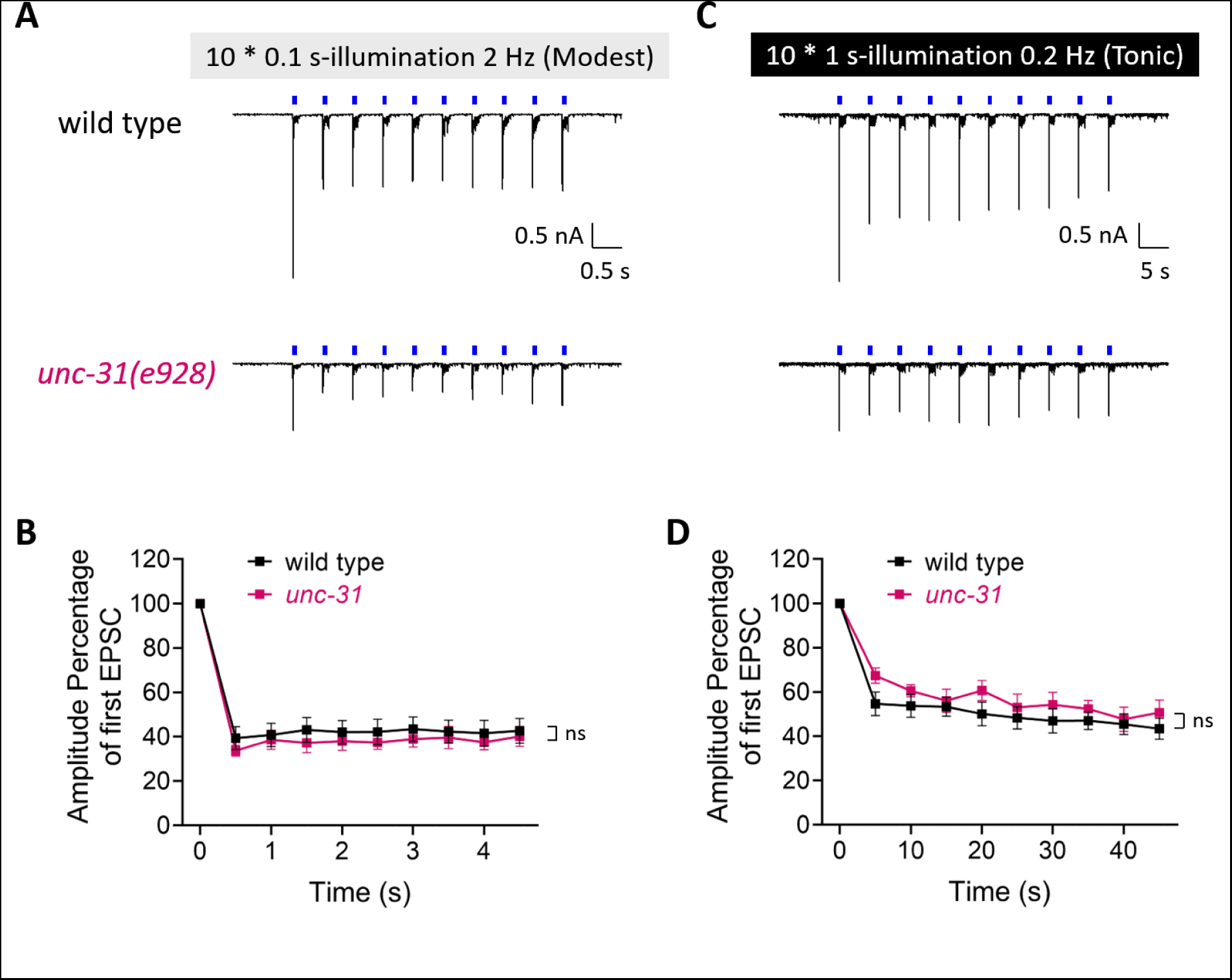
UNC-31 is not required for stimulation induced suppression of EPSCs. (A, C) Sample traces of sequential evoked EPSCs elicited by 10 consecutive 0.1 s blue light pulses at 2 Hz (A), or 10 consecutive 1 s blue light pulses at 0.2 Hz (C) recorded from WT and *unc-31(e928)* worms. (B, D) Quantification of eEPSCs amplitude normalized relative to the first evoked current for the indicated strains. Differences in traces were analyzed by two-way ANOVA. ns, no significance.

